# Numb is required for optimal contraction of skeletal muscle

**DOI:** 10.1101/2021.09.03.453960

**Authors:** Rita De Gasperi, Chenglin Mo, Daniella Azulai, Zhiying Wang, Lauren M. Harlow, Yating Du, Zachary Graham, Jiangping Pan, Xin-hua Liu, Lei Guo, Bin Zhang, Fred Ko, Ashleigh M Raczkowski, William A. Bauman, Chris N Goulbourne, Wei Zhao, Marco Brotto, Christopher P. Cardozo

**Author notes:** Address Correspondence to: Christopher Cardozo, MD, Center for the Medical Consequences of Spinal Cord Injury, James J. Peters VA Medical Center, 130 West Kingsbridge Road, Bronx, NY 10468. Contributed equally to the design of the research and data interpretation (Rita De Gasperi, Chenglin Mo, Marco Brotto, Christopher P. Cardozo).

## Abstract

**Background:** The role of Numb, a protein that is important for cell fate and development was investigated in adult skeletal muscle in mice using a conditional, inducible knockout (cKO) model.

**Methods:** Numb expression was evaluated by Western blot. Numb localization was determined by confocal microscopy. The effects of cKO of Numb and the closely-related gene Numb-like in skeletal muscle fibers was evaluated by *in-situ* physiology; transmission and focused ion beam scanning electron microscopy; 3-dimensional reconstruction of mitochondrial; lipidomics; and bulk RNA-sequencing. Additional studies using primary mouse myotubes investigated the effects the effects of Numb knockdown on cell fusion, mitochondrial function and calcium transients.

**Results:** Numb protein expression was reduced by ∼70% (p < 0.01) at 24 as compared to 3 months of age. Numb was localized within muscle fibers as bands traversing fibers at regularly spaced intervals in close proximity to dihydropyridine receptors.

The cKO of Numb and Numb-like reduced specific tetanic force by 36%, p < 0.01), altered mitochondrial spatial relationships to sarcomeric structures, increased Z-line spacing by 30% (p < 0.0001), perturbed sarcoplasmic reticulum organization and reduced mitochondrial volume by over 80% (p < 0.01). Only six genes were differentially expressed in cKO mice: *Itga4, Sema7a, Irgm2, Vezf1, Mib1* and *Tmem132a*. Several lipid mediators derived from polyunsaturated fatty acid (PUFAs) through lipoxygenases were upregulated in Numb cKO skeletal muscle; 12-HEPE was increased by ∼250% (p < 0.05) and 17,18-EpETE by ∼240% (p < 0.05). In mouse primary myotubes, Numb knock-down reduced cell fusion (∼20%, p < 0.01) and mitochondrial function and delayed the caffeine-induced rise in cytosolic calcium concentrations by more than 100% (p < 0.01).

**Conclusions:** These findings implicate Numb as a critical factor in skeletal muscle structure and function which appear to be critical for calcium release; we therefore speculate Numb plays critical roles in excitation-contraction coupling, one of the putative targets of aged skeletal muscles. These findings provide new insights into the molecular underpinnings of the loss of muscle function observed with sarcopenia.

## INTRODUCTION

Aging is associated with sarcopenia, a condition associated with deleterious changes in skeletal muscle that include progressive loss of muscle mass and reduced muscle quality reflected as impaired force generation and power [1]. Molecular mechanisms responsible for age-related loss of muscle quality remain incompletely understood. Mounting evidence supports a role for decreased effectiveness of excitation-contraction coupling whereby the multiprotein complex called the triad couples membrane depolarization with calcium release. [2]. Deficiency in calcium regulatory components contributes greatly to reduced muscle function during aging [3-5]. Reduced store-operated calcium entry (SOCE) function [6, 7] has also been implicated.

Altered balance in the generation of pro-inflammatory lipid mediators (LMs) and anti-inflammatory LMs during aging leads to low-grade systemic inflammation may also contribute to the development of various age-related disorders such as sarcopenia [8].

Determinants of calcium release during excitation-contraction coupling remain incompletely understood. Studies in cell culture suggest that one regulator of calcium release is the adaptor protein Numb [9]. Numb mRNA expression was reduced in muscle of older individuals [10] raising questions regarding the role of reduced Numb expression in the pathogenesis of aging-related decline of muscle strength and power. Numb is a highly conserved protein expressed in all higher organisms [11]. A closely related gene, Numb-like (NumbL) has also been identified [11]. NumbL function overlaps with though is distinct from that of Numb [12]. Numb has roles in development, cell fate commitment, termination of Notch and Sonic Hedgehog (Shh) signaling [13, 14], mitochondrial fission in kidney [15] and plays an important role in skeletal muscle satellite cells [16].

The possibility that Numb contributes to function of adult skeletal muscle fibers has not been previously investigated. Here we report that, in adult mice, Numb is concentrated within skeletal muscle fibers where its expression is required for optimal muscle force production, sarcomere structure and morphology of intermyofibrillar mitochondria. Studies with primary cultures of mouse myotubes link Numb to myoblast fusion, mitochondrial function, and calcium release from intracellular stores. Underlying mechanisms were explored using RNA-sequencing and lipidomics profiling focused on signaling lipids formed from n-3 and n-6 unsaturated fatty acids.

## Methods

### Animals

C57B6N mice used for determination of changes in Numb protein levels were obtained from the NIA Aged Rodent Colonies and euthanized at the ages indicated. C57BL/6 mice used for immunofluorescence studies were from Charles River Laboratories. Mice carrying a floxed allele for the Numb (fl-Numb) and Numb-like (fl-NumbL) genes [18], were obtained from Jackson Laboratories (Bar Harbor, ME) *(Numb*^*tm1Zili*^ *NumbL*^*tm1Zili*^/J, stock # 005384). The fl-Numb and fl-NumbL lines were heterozygous for one or both alleles at the time they were received and were crossed until homozygous for both floxed Numb and floxed NumbL; these animals are designated Numb^f/f^/NumbL^f/f^. HSA-MCM mice, which express a mutated-estrogen receptor-Cre recombinase (Mer-Cre-Mer) double fusion protein under the control of the human actin alpha-1 (skeletal muscle) promoter [17] were a generous gift from Dr. Karyn Esser, University of Florida. Mice were genotyped using genomic DNA isolated from tail or ear snips. The primers for the floxed Numb allele were as described [18] while the following primers were used for the floxed NumbL allele: 5’ GAGTTTCCGTACATGCTTTGGG and 5’ GGAGACCTTCTCAATGGTCTGG.

Cre genotyping was performed as previously described [38]. The study protocol was approved by the James J Peters VA Medical Center Institutional Animal Care and Use Committee. All animal procedures were conducted in accordance with the requirements of Guide for the Care and Animal Use of Laboratory Animals.

### Tamoxifen treatment

HSA-MCM Numb^f/f^/NumbL^fl/f^ mice were injected intraperitoneally with 0.2 ml of tamoxifen solution (10 mg/ml in peanut oil/5% ethanol) or vehicle for 5 consecutive days and weekly thereafter beginning on day 10 after the first injection until they were sacrificed at 56 days after the first injection. These mice are referred to as Numb/NumbL cKO and vehicle-treated controls, respectively. cKO, Numb^f/f^/NumbL^fl/f^ mice treated with tamoxifen or vehicle were used as tamoxifen-treated or vehicle-treated genotype controls, respectively.

### Statistical analysis

Data are shown as mean value ± standard deviation (SD). Data were analyzed by ANOVA with post-hoc Tukey HSD test or by a two-tailed unpaired t-test. The statistical analyses were performed with OriginPro 2020 (Figures 5-7 and Supplemental Figure S12) or Graph Pad Prism 9.0 (all other figures). A value for p<0.05 was considered significant.

## RESULTS

### Numb and NumbL expression and localization in muscle

Numb expression was investigated in skeletal muscle by western blot using gastrocnemius muscles isolated from 3 and 24-month-old C57BL/6N. These data showed that Numb was expressed in muscle but significantly lower levels of Numb protein were found in 24 months old mice when compared to 3 months old mice (Supplementary Fig.1), indicating age-dependent decreased expression of Numb protein in mouse skeletal muscle.

NumbL expression could be detected by qPCR and was approximately 5-10 fold lower than that of Numb (Supplementary Fig.2).

To determine the localization of Numb in skeletal muscle, sections of mouse gastrocnemius muscle from C57B6 mice were immunostained for Numb and dystrophin. Intense Numb staining was observed within myofibers where it was organized in distinct bands that traversed the myofiber at regular intervals (Fig 1a-c). A similar Numb distribution was observed by immunostaining of muscle fibers isolated from mouse hindlimb muscles (Fig. 1d). Numb immunostaining did not colocalize with the Z-line protein α-actinin but showed a precisely ordered relationship to it indicating a spatial relationship of Numb to the sarcomere (Fig. 1e).

**Fig. 1.**
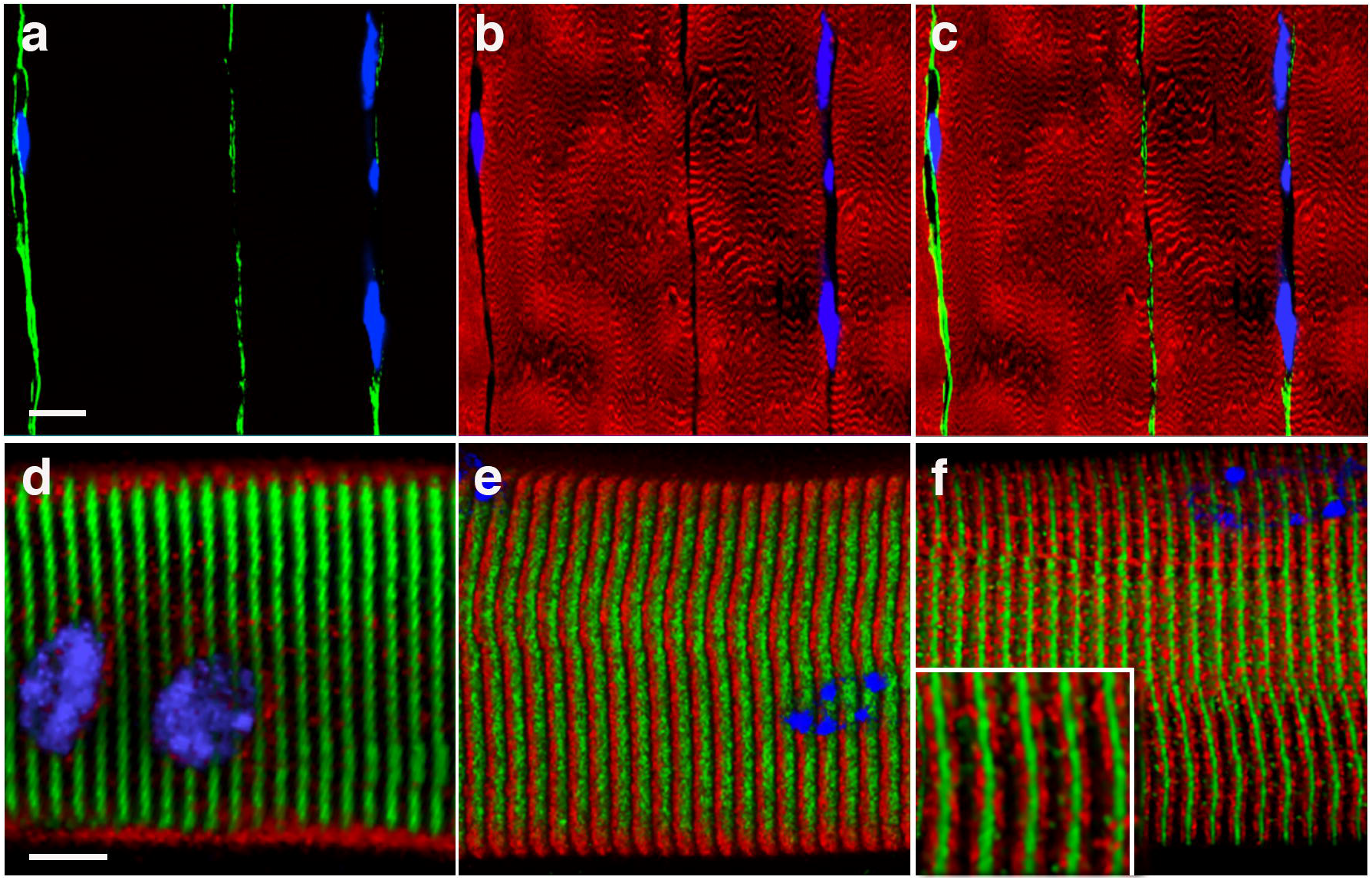
Numb protein localizes to the sarcomere of skeletal muscles. (a-c) Representative longitudinal sections of mouse gastrocnemius muscle immunostained for dystrophin (green, a, and Numb (red, b); the merged image is shown in panel (c). (d-f) Z-stacks of confocal images of isolated muscle fibers stained for Numb (green) and with: dystrophin(d); actinin(**e**) and dihydropyridine receptor (DHPR) (f). The insert in (f) shows a higher magnification of the Numb/DHPR stained fiber. Nuclei (blue) are labeled with DAPI in every panel. Scale bar: 10 μm; 5 μm for the insert in (f).

Numb appeared to be in close proximity to, though not overlapping with, the dihydropyridine receptor (DHPR) (Fig. 1f).

### Generation and characteristics of muscle specific inducible Numb conditional knock-out

To understand the physiological role(s) of Numb in skeletal muscle fibers, mice in which Numb expression was conditionally and inducibly knocked out in muscle fibers were generated.

Because NumbL may substitute for Numb in some biological contexts and might be upregulated after knocking out Numb, the conditional, inducible knockout was designed to inactivate both genes. Mice expressing a chimeric Cre recombinase sandwiched between two copies of a mutated estrogen receptor ligand binding domain (MCM) under the control of a fragment of the human skeletal actin promoter [17] (HSA-MCM) were crossed with mice homozygous for floxed Numb and NumbL alleles (Numb^f/f^/NumbL^f/f^) [18]. The resulting HSA-MCM Numb^f/f^/NumbL^f/f^ mice were treated with tamoxifen (Numb/NumbL cKO) or vehicle for 5 days and then once weekly until the time of euthanasia at 56 days. Numb protein levels were reduced in gastrocnemius (GS), tibialis anterior (TA) and soleus muscle (Fig. 2 a-b) after tamoxifen treatment as compared to vehicle treated mice.

**Fig. 2.**
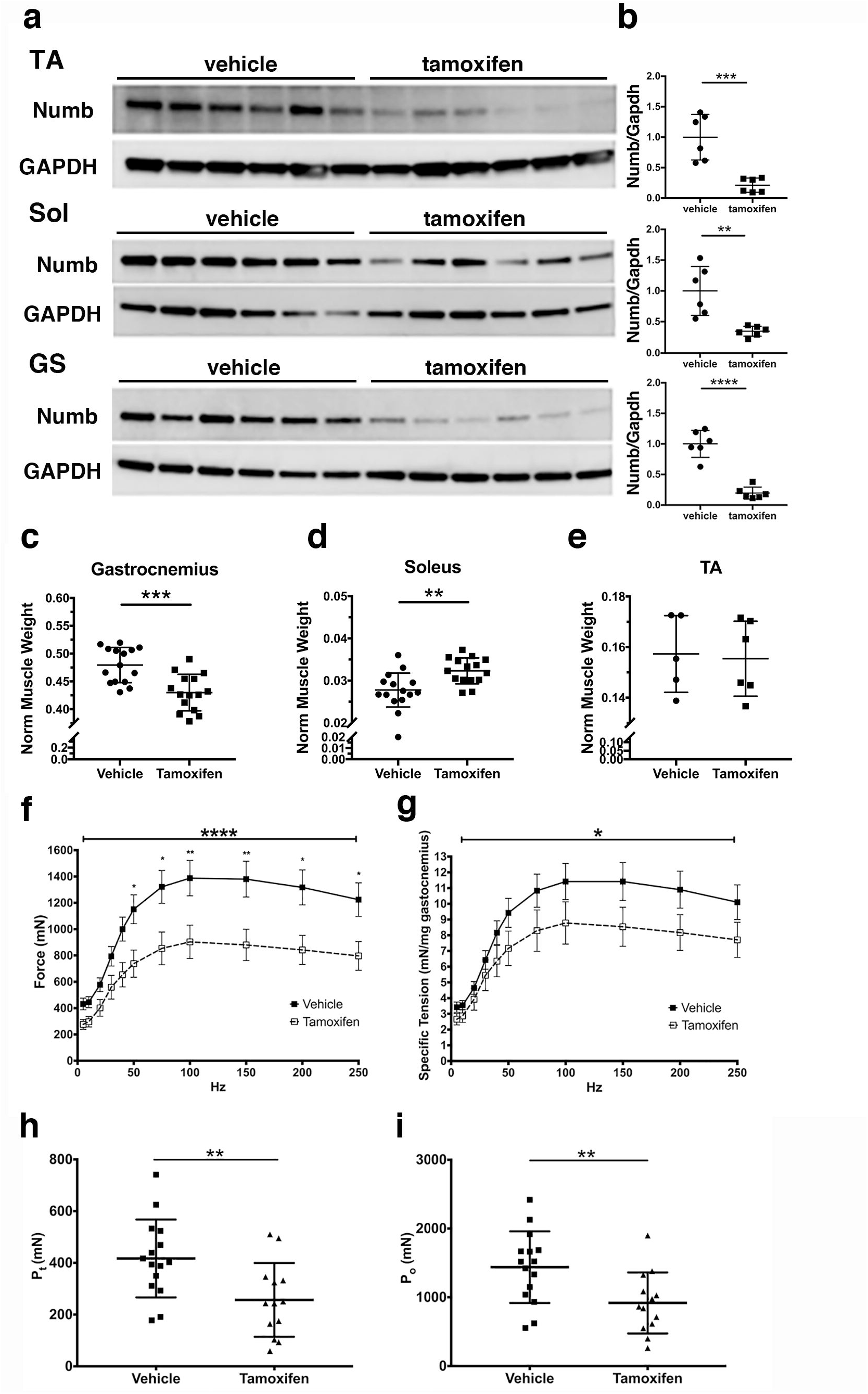
Numb/NumbL specific knock down in skeletal muscle leads to drastic decrease in contractile muscle force. HSA-MCM Numb^fl/fl^/NumbL^fl/fl^ mice were treated with vehicle or tamoxifen and sacrificed 56 days post-induction. (a) Western blot analysis showing reduced levels of Numb protein in Tibialis anterior (TA), Soleus (Sol) and Gastrocnemius (GS) muscles (N=6/group). (b) Quantitation of blots shown in (**a**). Numb levels were normalized to GAPDH (loading control) and expressed relative to the mean level of Numb in vehicle-treated samples. (N= 6/group). (c-e) Normalized muscle weights; wet muscle weight at time of euthanasia was normalized to pre-induction body weight. (N=15/group). (f-i) Force-frequency (f) and specific force frequency (g) relationships during *in-situ* physiological testing of the gastrocnemius muscle; (h) peak twitch force (Pt); (**i**) tetanic force (Po); N=15 for vehicle and 13 for tamoxifen-treated mice. Statistical analysis was performed with repeated measure one-way ANOVA with post-hoc Tukey HSD test (f-g) (Force frequency, F=4.753, DFn 10, DFd 260, Drug x Frequency interaction p<0.0001; specific tension force frequency, F=2.150, DFn 10, DFd260, Drug x Frequency interaction p=0.0213) or two-tailed unpaired t-tests (all other panels). **** p<0.0001, ***p< 0.001, ** p<0.01, * p< 0.05.

Body weights of Numb/NumbL cKO mice were not significantly different from vehicle-treated controls and tamoxifen did not alter body weights of genotype-controls (Numb^f/f^/Numb^f/f^; Supplementary Fig. S3 a, f). Individual muscle weights of Numb/NumbL cKO showed a small decrease for gastrocnemius (Fig. 2c), plantaris, biceps and triceps (Supplementary Fig. S3 b,d,e), while no significant changes were observed for TA (Fig. 2 e) or EDL (Supplementary Fig. S3c). Soleus showed a small increase in weight (Fig. 2 d), possibly representing compensatory hypertrophy due to weakening of gastrocnemius and plantaris. Administration of tamoxifen to genotype control Numb^f/f^/NumbL^f/f^ mice did not alter muscle weights, with the exception of a small decrease in plantaris weight (Supplementary Fig.S3 g-l).

### Numb/NumbL cKO alters muscle contractile properties

The effect of Numb/NumbL cKO on gastrocnemius muscle contractile properties was evaluated by *in-situ* physiology. Compared to vehicle treated animals, Numb/NumbL cKO mice demonstrated significantly reduced absolute and specific force (mN/mg muscle), peak twitch and peak tetanic force (Fig. 2 f-i), while muscle fatigue, time to peak tension and half-relaxation time were not significantly altered (Supplementary Fig. S5 a-c). No difference was observed between Numb/NumbL cKO male and female mice (Supplementary Fig. S4). Administration of tamoxifen to Numb^f/f^/Numb^f/f^ genotype controls did not alter muscle contractile properties based on analysis of force-frequency curves (Supplementary Fig. S6).

### Numb/NumbL cKO did not alter muscle cross-sectional area (CSA) or fiber types

Analysis of CSA of TA muscle showed no significant difference between Numb/NumbL cKO animals and vehicle-treated mice (Supplemental Fig. S7 a-c). Analysis of fiber-type composition of TA muscles with antibodies against myosin heavy chain isoforms did not show any significant difference between Numb/NumbL cKO mice and vehicle injected mice (Supplementary Fig. S7 d-h).

To exclude any tamoxifen effect that was independent of the Numb/NumbL knockout, muscle fiber CSA, distribution and type were determined in TA muscles derived from Numb^f/f^/NumbL^f/f^ mice treated as above. CSA was slightly (∼3%) though significantly increased in tamoxifen treated mice as compared to vehicle-treated controls (Supplementary Fig. S8 a-c), while fiber type composition was not altered by tamoxifen administration (Supplementary Fig. S8 d-e).

### Numb/NumbL cKO Perturbs Muscle Ultrastructure

To gain insights as to why muscle force production was reduced by Numb/NumbL cKO, muscle ultrastructure was examined in TA muscle by transmission electron microscopy. This analysis revealed multiple ultrastructural alterations in Numb/NumbL cKO (Fig.3b) as compared to control (Fig 3a). Frequent discontinuities of Z-disks were observed (Fig. 3b, Arrow 1).

**Fig. 3.**
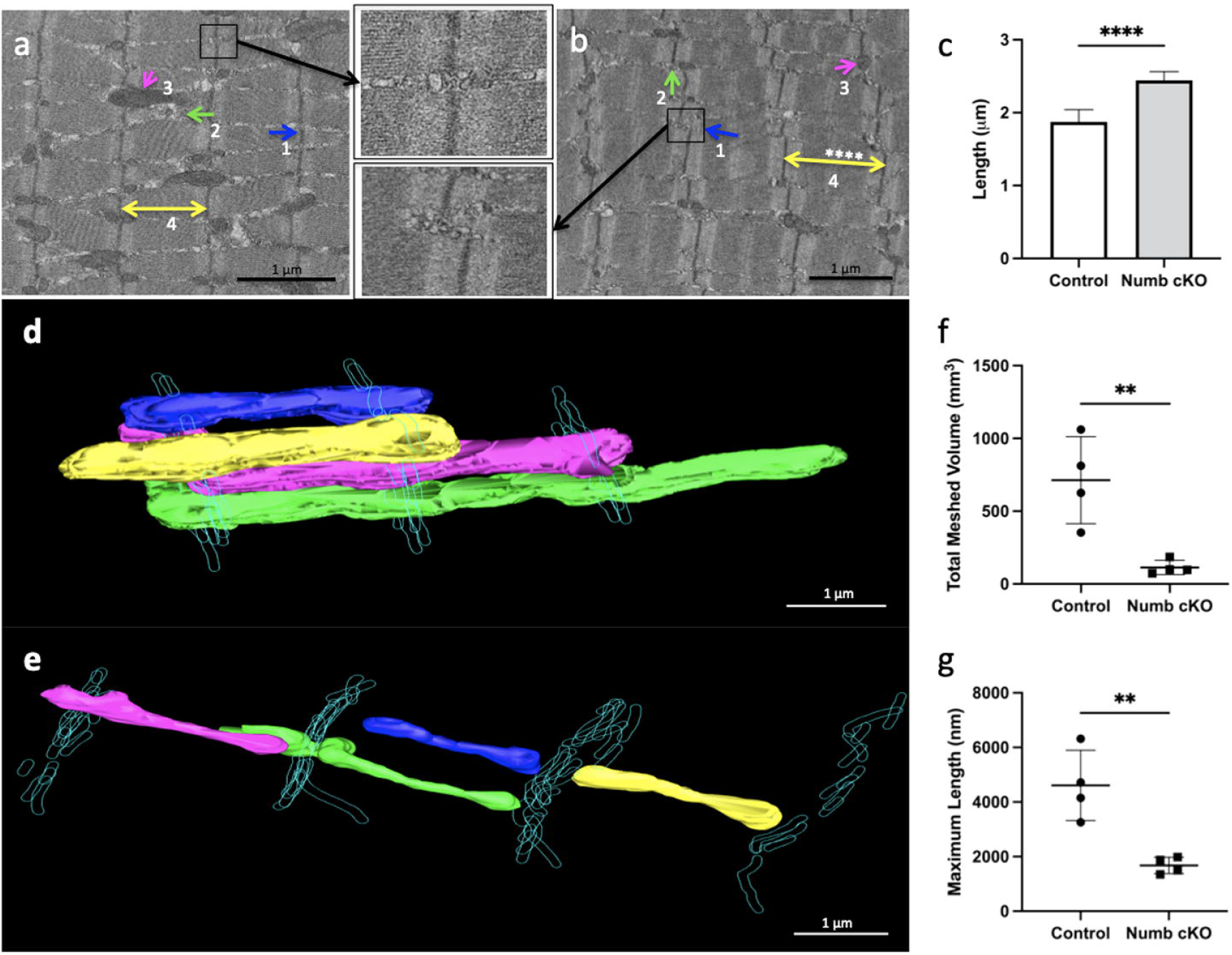
Numb knockout resulted in marked ultrastructural changes in skeletal muscle. Transmission electron microscopy was performed on longitudinal sections of mouse tibialis anterior muscle at 5,300x. Representative images for HSA-MCM/Numb^f/f^/NumbL^f/f^ mice treated with vehicle (a) or tamoxifen (b) are shown. Arrows indicate (1) the positions of Z-lines, (2) sarcoplasmic reticulum, (3) intermyofibrillar mitochondria, (4) Z-disc spacing. (c) Graph depicts the distance between Z-discs (m) for three animals per group. **** p < 0.0001, two-tailed unpaired t-test. Shown between (a) and (b) are enlarged images of selected areas (box in panel a-b). (d-e) Three-dimensional reconstruction of mitochondria in TA muscle of control (d) and Numb/NumbL cKO mice (e). Shapes outlined in blue represent Z-discs; solid shapes represent individual mitochondria. (f-g) Maximum length (f) and meshed volume (g) of mitochondria in (d) and (e). Scale bars, 1 μ m.

Mitochondria were smaller and had lost their normal spatial organization relative to the sarcomere (Fig. 3b, Arrow 2). Sarcoplasmic reticulum was disorganized with increased spacing between the linear clusters dividing the contractile elements on longitudinal sections (Fig. 3b, Arrow 3). Distances between Z-lines were longer in TA muscle from Numb/NumbL cKO mice (Fig. 3b, Arrow 4 and Fig. 3c). Tamoxifen had no effect on the ultrastructure of TA muscle from Numb^f/f^/NumbL^f/f^ mice (Supplementary Fig. S9).

To more closely examine changes in mitochondria, TA muscle from Numb/NumbL cKO and control were examined by focused ion beam scanning electron microscopy. Residual bodies were frequently observed in muscle from Numb/NumbL cKO mice (Supplementary Fig. S10) but not in controls. Three-dimensional reconstruction of mitochondria showed that mitochondria in Numb/NumbL cKO mice were smaller than control (f), typically did not cross the Z-disk, and had less complex shapes (Fig. 3e; Supplementary Materials Movies 1 and 2). Quantitation of the volume and length of mitochondria showed that both volume and length were reduced in Numb/NumbL cKO muscle (Fig. 3 f-g).

### Numb/NumbL cKO alters the expression of a very select group of genes

To dissect potential molecular mechanisms by which Numb/NumbL cKO regulates muscle function, RNA sequencing was performed using RNA from gastrocnemius muscle from tamoxifen or vehicle treated HSA-MCM/Numb^f/f^/NumbL^f/f^ male and female mice. RNA from gastrocnemius from male and female Numb^f/f^/NumbL^f/f^ (control genotype) mice treated as above was also sequenced to serve as a drug treatment control. Four hundred and seven differentially expressed genes (DEGs) were identified in female HSA-MCM/Numb^f/f^/NumbL^f/f^ muscle after tamoxifen treatment (Supplementary Table 1), while only 35 DEGs were found in tamoxifen-treated females of the control genotype (Supplementary Table 2). In male Numb/NumbL cKO mice, 187 genes were differentially regulated (Supplementary Table 3), while 705 DEG were observed in tamoxifen treated male genotype control mice (Supplementary Table 4 and Supplementary Figure S11). After removing those genes altered by tamoxifen in genotype controls, 389 DEGs were identified in female Numb/NumbL cKO muscle (Fig 4a), (212 upregulated and 177 downregulated), while only 68 DEG were identified in males (Figure 4b) (39 downregulated and 29 upregulated).

**Fig 4.**
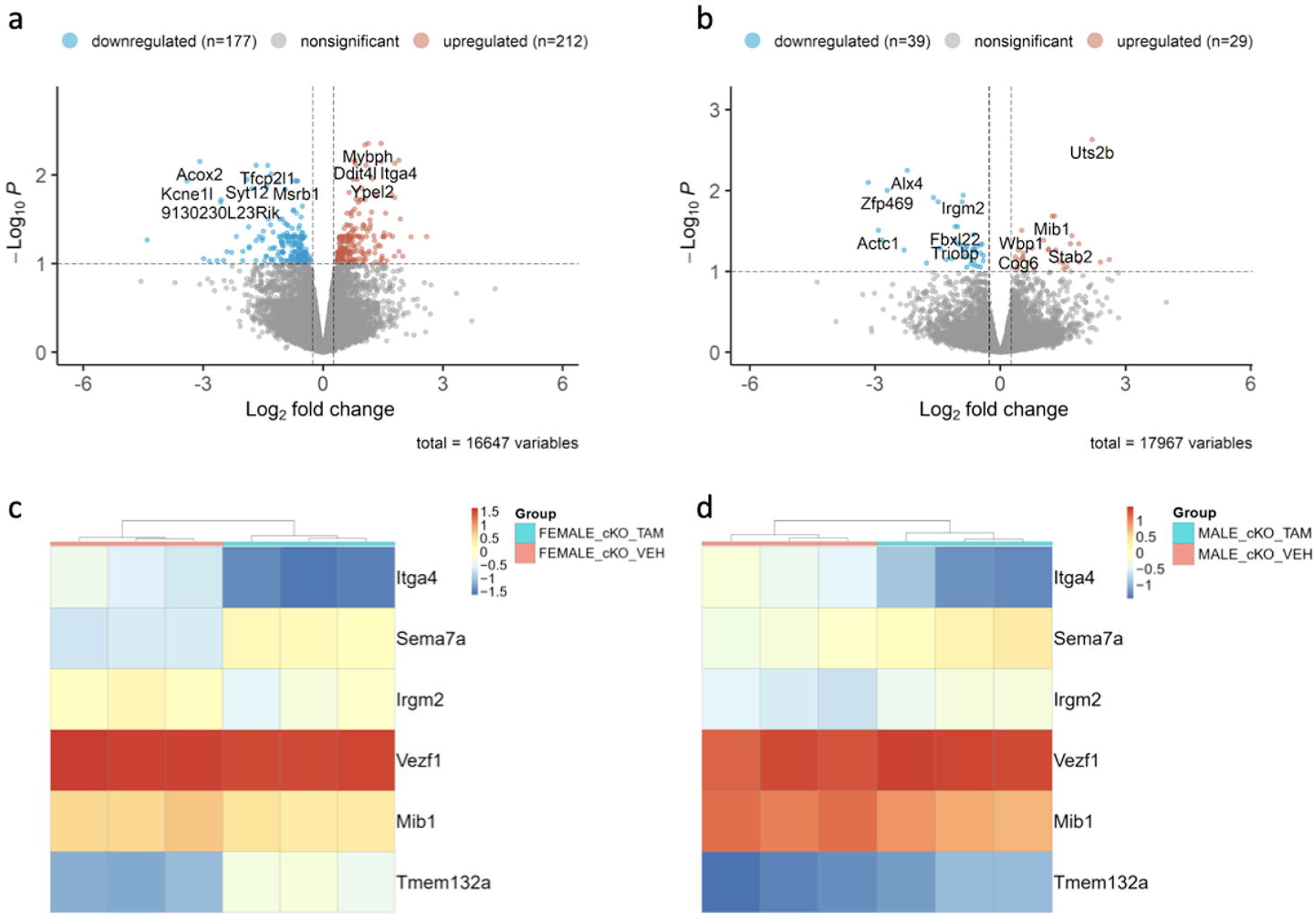
RNA sequencing identified genes that are differentially expressed after Numb/NumbL cKO. (a-b) Volcano plots showing mRNA expression levels for genes with fold-changes greater than 1.2 and p-values adjusted for FDR of less than 0.1 (a, females, b males). Differentially expressed genes are shown as light brown spots (upregulated) or light blue spots (downregulated). All other genes detected are shown as grey dots. Shown at the top of each panel are the values for numbers of up and downregulated genes after removal of genes altered in tamoxifen treated genotype control females and males mice respectively. The total number of transcripts analyzed is shown under the X-Axis as total variables; (c-d) Heatmaps showing relative expression of genes altered in both female (c) and male (d) Numb/NumbL cKO mice.

Six genes were differentially regulated in both male and female Numb/NumbL cKO (Figures 4c, d and Supplementary Table 5). Heatmaps showed that expression patterns for each of these genes were similar within each group (Figs. 4c, d). Two genes were downregulated in both males and females: Sema7a, a cytoplasmic membrane protein involved in integrin signaling and Tmem132a, a gene with poorly understood function (Figs. 4c, d). Three genes were upregulated in both males and females: Itga4, an α-integrin subunit; Mib1, an E3 ubiquitin ligase that promotes internalization and activation of Notch receptors and VezF1, a transcription factor that may be involved in angiogenesis and development (Figures 4c, d). Irgm2, a GTPase implicated in innate immune function, was upregulated in females but downregulated in males (Figures 4c, d).

### Alterations in lipid mediator levels occur after Numb/NumbL cKO

The Lipid Signaling Mediator (LM) profiles of gastrocnemius muscles were measured using an LC-MS/MS-based method. Levels for 158 LMs, primarily derived from omega-6 and omega-3 polyunsaturated fatty acids including arachidonic acid (AA), eicosapentaenoic acid (EPA), and docosahexaenoic acid (DHA), were determined. Forty-eight lipid mediators were altered in gastrocnemius muscles from Numb^f/f^/NumbL^f/f^ and HSA-MCM/ Numb^f/f^/NumbL^f/f^ mice treated with vehicle or tamoxifen for 56 days. The percentage change of each lipid mediator was calculated by dividing its relative area [(peak area of analyte/peak area of Internal Standard) in LC chromatogram] to that in vehicle-treated Numb^f/f^/NumbL^f/f^ mice. The relative levels in Numb/NumbL cKO mice for selected key lipid mediators are shown in Fig. 5 (a-f), and the list of all 48 identified LMs has been summarized in Supplementary Table 6. Tamoxifen altered levels of some LM in Numb^f^/NumbL^f/f^ cKO mice without altering LM levels in Numb^f/f^/NumbL^f/f^ genotype control mice (Fig. 5a-f). Specifically, in Numb/NumbL cKO mice, there was a significant increase in the levels of 12-hydroxyeicosapentaenoic acid (12-HEPE, p=0.035), 17,18-epoxyeicosatetraenoic acid (17,18-EpETE, p=0.041), 12-hydroxyeicosatetraenoic acid (12-HETE, p=0.024), leukotriene B_4_ (LTB_4_, p=0.090), 13-oxo-9,11-octadecadienoic acid (13-KODE, p=0.032), and N-arachidonoylethanolamine (AEA, p=0.032). Of note, 12-HEPE, 12-HETE, LTB_4_, and 13-KODE are derived from arachidonic acid (AA), linoleic acid (LA), and eicosapentaenoic acid (EPA), respectively, *via* lipoxygenase (LOX) mediated pathways (Fig. 5g). These data suggest that the observed phenotypes of Numb/NumbL cKO mice could be associated with the biological activities of these LMs and, more broadly, with activation of LOXs and their downstream pathways.

**Fig. 5.**
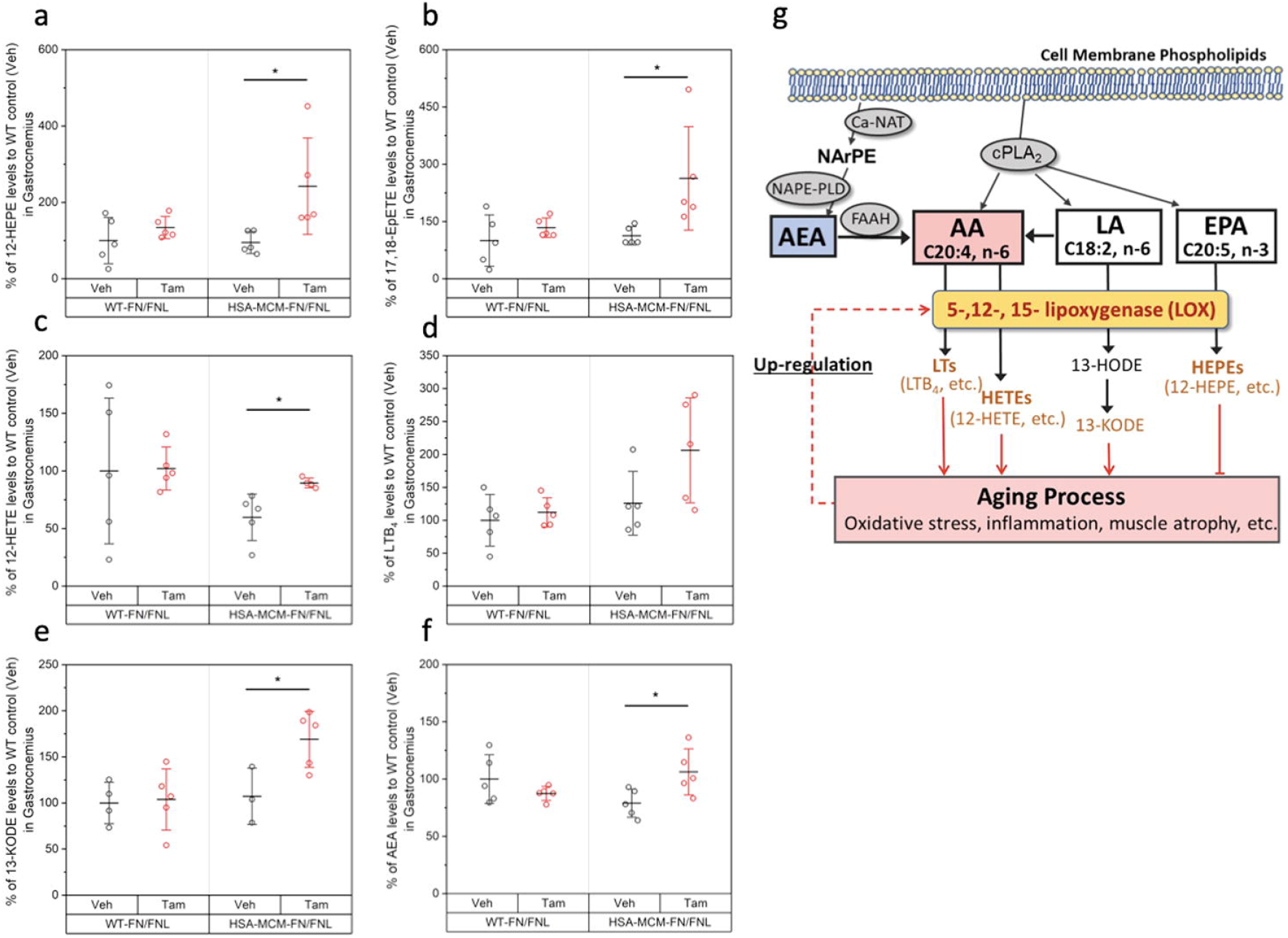
Numb/NumbL KO leads to significant alterations in the levels of key lipid signaling mediators associated with the lipoxygenase pathway in gastrocnemius muscles. (a) 12-HEPE; (b) 17,18-EpETE; (c) 12-HETE; (d) LTB_4_; (e) 13-KODE; (f) AEA. Data are shown as mean ± SD, n=5 for each group, two-tailed Student’s t-test was applied for mean comparison with Vehicle (Veh), *p<0.05. WT-FN/FNL: Numb^f/f^/NumbL^f/f^; HSA-MCM-FN/FNL: HSA-MCM/ Numb^f/f^/NumbL^f/f^; Tam, tamoxifen; Veh, vehicle; (g) The most altered signaling LMs in Numb/NumbL cKO mice are derived from the 5-, 12- or 15-lipoxygenase (LOX) pathways. NArPE, N-arachidonoyl phosphatidylethanolamine; AEA, N-arachidonoylethanolamine; AA, arachidonic acid; LA, linoleic acid; EPA, eicosapentaenoic acid; LTs, leukotrienes; HETEs, hydroxyeicosatetraenoic acids; 13-HODE, 13-hydroxyoctadecadienoic acid; 13-KODE, 13-oxo-9,11-octadecadienoic acid; HEPEs, hydroxyeicosapentaenoic acids; Ca-NAT, Ca^2+^-dependent N-acyltransferase; NAPE-PLD, N-acyl-phosphatidylethanolamine-specific phospholipase D; FAAH, fatty acid amide hydrolase. AEA is biosynthesized from membrane phospholipids through the action of a Ca^2+^-dependent N-acyltransferase (Ca-NAT) and then of N-acyl-phosphatidylethanolamine-specific phospholipase D (NAPE-PLD). AEA is subsequently degraded by fatty acid amide hydrolase (FAAH) to generate AA.

### Acute Numb KO reduces fusion of mouse primary myoblasts *in vitro*

To further investigate the mechanism of Numb on the functions of skeletal muscle, mouse primary myoblasts were treated with a vivo-morpholino against Numb. Treatment with Numb morpholino for 48 h led to approximately 30% reduction in Numb protein levels (Supplementary Fig. S12a-b) and 15% reduction in fusion index (Supplementary Fig. S12c-d), suggesting that Numb plays an important role in myoblast fusion.

### Acute Numb knockdown Reduces Mitochondrial Functional Capacity of Mouse Primary Myoblasts

To specifically determine whether the physiological and ultrastructural changes in the cKO mice could be associated with mitochondrial dysfunction, mitochondrial function was compared between control and Numb morpholino-treated primary mouse myotubes using the Seahorse energy phenotype test kit and mitochondrial stress test kit. Compared with control, the Numb knockdown group had lower mitochondrial and glycolytic energy production although the balance between the two was preserved (Fig. 6a). Following the phenotype test, the mitochondrial stress test was used to determine the effect of Numb knockdown on mitochondrial respiration. As shown in Fig. 6d-f, Numb knockdown reduced basal respiration, maximal respiration, and spare capacity but did not alter non-mitochondrial oxygen consumption (Fig. 6c), or ATP-linked respiration (Fig. 6g). In Numb-knockdown myotubes, oxygen consumption rate (Fig. 6h) was decreased both at baseline and when mitochondria were stressed, while extracellular acidification rate was decreased only in stressed condition (Fig. 6i).

**Fig. 6.**
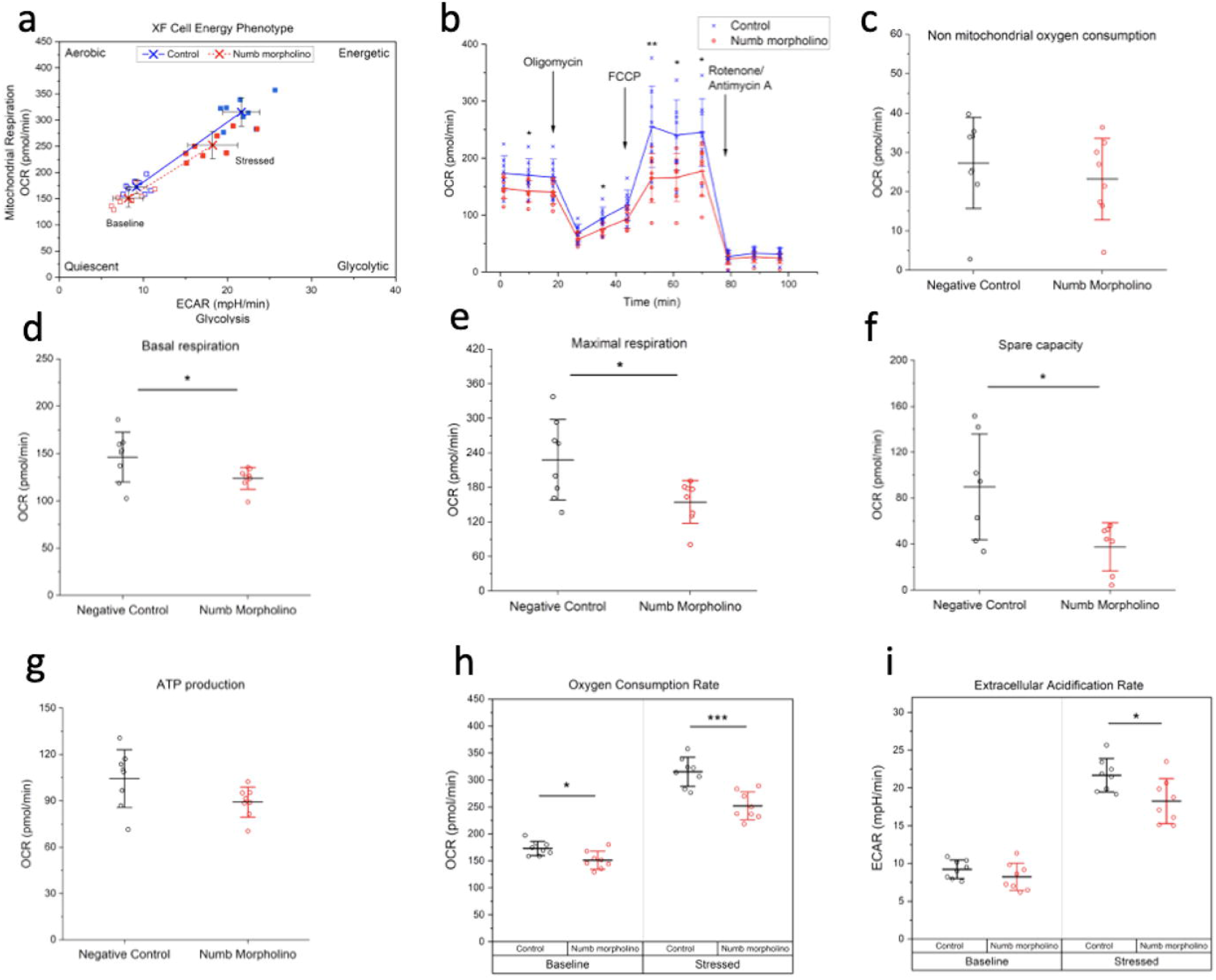
Acute knockdown of Numb negatively impacted mitochondrial function. (a) Less energetic phenotype resulted from knockdown of Numb; (b) overall profiling of mitochondrial functions determined by a Seahorse mitochondrial stress kit; (c) Non mitochondrial oxygen consumption; (d) basal respiration; (e) maximal respiration; (f) spare capacity; (g) ATP production; (h) oxygen consumption rate; (i) extracellular acidification rate. The data shown are representative results from four independent experiments. Data are shown as mean ± STD. *, p < 0.05; **, p < 0.01.

### Effect of Numb on cytosolic calcium transients

Due to the importance of intracellular calcium homeostasis to the control of muscle function, and the prominent role of excitation-contraction coupling in muscle contractile force generation, one explanation for some of our findings in the Numb/NumbL cKO mice would be a decreased release of calcium ions from sarcoplasmic reticulum stores during excitation-contraction coupling. To assess the role of Numb in cytosolic calcium homeostasis, caffeine-induced calcium transients were measured in cultured mouse primary myotubes treated with a Numb targeted vivo-morpholino to downregulate this gene. Control cells were treated with a biologically inactive, control vivo-morpholino. In cells treated with the control morpholino, the baseline Fura-2 signal was stable; addition of 20 mM caffeine to cultures stimulated a robust increase in fluorescence (Fig. 7a). In contrast, in myotubes treated with a Numb-targeting vivo-morpholino, the development of spontaneous calcium release events was observed as small calcium oscillations as if calcium induced calcium release (CICR) was overactive in these fibers. Moreover, the caffeine-induced calcium release transient observed in Numb-knockdown cells appeared to be reduced by more than 50% (Fig. 7a) and the time taken for responding to caffeine stimulation (time to peak) was significantly prolonged by Numb knockdown (Fig. 7a-b).

**Fig. 7.**
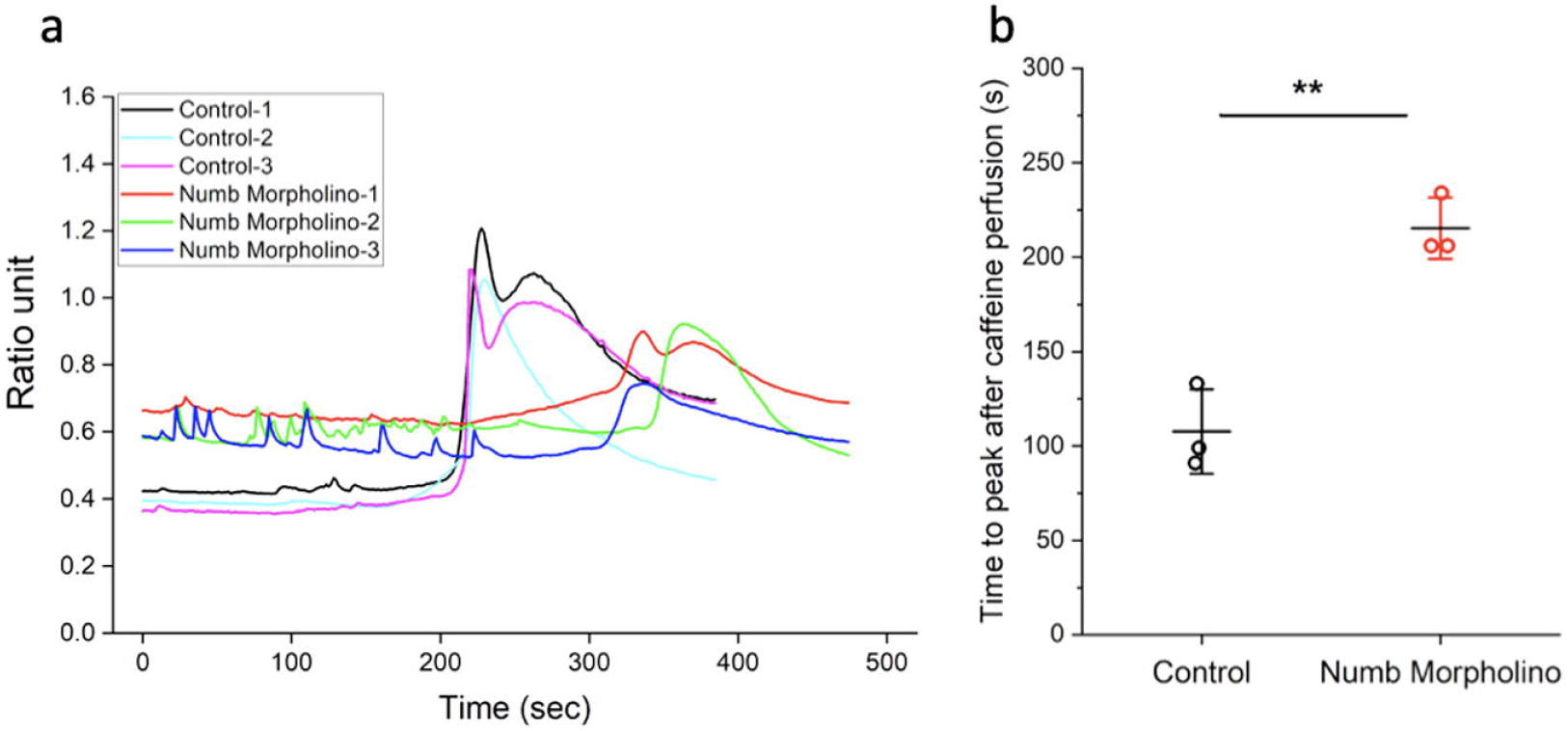
Numb KO elicits spontaneous CICR events and reduces caffeine-induced SR Ca^2+^ release. Primary myoblasts were treated with either a control or Numb-targeting vivo-morpholino. The cells were loaded with Fura-2 AM and intracellular calcium transients induced by 20 mM caffeine were monitored based on F350/F375 readings. (a) Representative tracings of F350/F375 over time for individual wells of cells treated with either Numb morpholino or control morpholino. (b) Time to peak F350/F375 was measured. 20-30 myotubes were tested for each group. Two-tailed unpaired Student’s t-test was applied for comparison with the Control, **p<0.01. Data are from 3-4 independent experiments.

## Discussion

The major conclusions supported by our study are that loss of Numb from skeletal muscle fibers results in significant loss of the capacity of muscle to generate force, and that Numb protein levels decline with advancing age in mouse gastrocnemius muscle. When coupled with prior observations that levels of Numb mRNA were reduced in muscle biopsy samples taken from men 60-75 years old [10], these data strongly support a causal linkage between reductions in Numb protein levels in skeletal muscle and sarcopenia-related loss of muscle force generating capability. While a role for NumbL in the physiological and ultrastructural phenotype observed in the Numb/NumbL cKO mice cannot be excluded, it seems unlikely that NumbL participates in function or homeostasis of healthy myofibers given the comparatively low expression of NumbL mRNA in adult skeletal muscle. This conclusion would be consistent with prior reports that mice carrying a loss-of-function mutation of NumbL had no appreciable muscle phenotype and that the mutation did not alter tissue repair [16]. Based on these observations, we propose that the physiological deficits observed in the Numb/NumbL knockout mice studied herein resulted essentially from decreased levels of Numb in muscle fibers.

Immunohistochemistry localized Numb close to DHPR, a component of the triad suggesting spatial and functional relationships between Numb and the function of the triad in the release calcium from SR stores as part of excitation-contraction coupling. Some support for this interpretations stems from the finding that knockdown of Numb in primary mouse myotubes reduced release of calcium from intracellular stores as determined by the caffeine-induced calcium release assay. The observation that in mouse primary myotubes, Numb knockdown results in spontaneous and transient rises in cytosolic calcium that is intriguingly similar to self-pacing calcium transients in beating cardiac myocytes, suggest that Numb knockdown could alter the regulation of CICR via triads, which is normally tightly controlled by membrane potential, therefore under the control of voltage and depolarization and not normally triggered as in the heart [19, 20].

Our data do not provide information as to the mechanism by which Numb regulates appropriate calcium release from the SR, but the localization of Numb near the DHPR supports several possibilities. Numb may bind to one or more triad proteins to change their capacity to release calcium from SR stores; alternatively, the altered mitochondrial function observed in myotubes in which Numb expression was reduced may increase release of reactive oxygen species which are deleterious to ryanodine receptor function [3]. Since caffeine acts via CICR to release calcium from the SR and enhanced spontaneous CICR was observed as if these muscles behaved like cardiac myocytes, it is also possible that the sensitivity of RyR-1 to CICR is significantly shifted to the left (i.e., super or hypersensitivity to calcium). In other words, it is possible that in these muscle cells, even at very low intracellular calcium levels, the RyR-1 is exquisitely sensitive to calcium, triggering CICR. It is possible that the spontaneous activity could lead to a partial depletion of the SR calcium stores, which in turn could result in the reduced overall response to caffeine. There are many other possibilities, such as altered ATPase, store-operate calcium entry (SOCE) function, as well as direct modifications within the contractile machinery, and an increased contribution of IP3 receptors to the oscillatory calcium pattern observed. These other possibilities were not studied here.

Lipid mediators are also regulators of calcium release from intracellular stores of C2C12 cell-derived myotubes [21] and smooth muscle [22]. Mo et al., demonstrated that nanomolar levels of PGE2 and agonists of its signaling pathway accelerate myogenic differentiation and proliferation of C2C12 muscle cells, primary mouse, and primary human muscle cells, while concomitantly causing intracellular calcium oscillations [21, 23, 8]. It is thus noteworthy that Numb knockdown upregulates concentrations of several LMs generated by LOXs and its downstream pathways. How these changes in LMs occur is not clear but it is noteworthy that increased intracellular calcium concentration such as might occur due to calcium leakage from intracellular calcium storages enhances the activities of LOXs and their upstream phospholipases resulting in increased production of LMs which could further exaggerate dysregulation of cytosolic calcium, and eventually contribute to the development of muscle weakness.

The elevations in LMs in muscle of mice with Numb/NumbL knockouts may have broader implications. Potent signaling lipids synthesized by oxidation of the PUFA precursors through COX and LOX are associated with various cellular process in the regulation of skeletal muscle mass and function [8, 23, 24]. Most ω-3 PUFA (EPA, DHA, etc.) metabolites are anti-inflammatory while those from ω-6 PUFA (e.g., AA and LA) are considered to be pro-inflammatory [25]. The enzyme 5/12/15-LOX plays a significant role in age-related muscle atrophy by different mechanisms ranging from regulating inflammation, protein degradation [26] and reactive oxygen species (ROS) generation [27]. In our lipidomics studies, the gastrocnemius muscles of Numb/NumbL knockout presented with significantly elevated levels for LTB_4_ and 12-HETE, which are pro-inflammatory eicosanoids derived from AA through the 5/12-LOX pathways. Similarly increased levels were also observed for 13-KODE, a potent ligand for PPAR-γ that is metabolized through the LA/13-HODE pathway which is associated with signaling of ROS production [28]. We speculate that subsequent exposure to these triggering factors (inflammation, ROS) leads to an abnormal chronic inflammatory condition, and ultimately contributes to aging-related muscle phenotypes in the Numb/NumbL cKO mice model. Additionally, another lipid mediator, 12-HEPE, was also significantly increased in skeletal muscles after Numb/NumbL depletion. 12-HEPE belongs to the ω-3 PUFA precursor EPA metabolic pathways, and is thought to attenuate inflammatory response in macrophages [29].

The increased 12-HEPE levels in the gastrocnemius of the Numb/NumbL cKO mice could be achieved by the up-stream lipoxygenase expression, therefore a likely compensatory adaptive mechanism during aging/inflammation. This hypothesis (summarized in Fig. 5g) is supported by the published observations that lipoxygenase is upregulated with aging [30]. All these findings suggest that lipoxygenase metabolic pathways play a functional role in the development of weakness due to Numb and NumbL knockouts and implicate lipoxygenase in sarcopenia.

Many studies have linked aging, mitochondrial dysfunction and sarcopenia [31, 32]. Reasons for the reduced size of mitochondria in Numb/NumbL cKO mice are unclear, but perhaps the accumulation of ROS and lipid damage could be a contributing factor. In addition, the accumulation of residual bodies in muscles of these mice suggest impaired quality control through mitochondrial fission and mitophagy. Analysis of cellular respiration of primary mouse myotubes depleted of Numb suggested reduced mitochondrial oxygen consumption rate, indicative of an overall decrease in mitochondrial respiration and ATP generating capacity. Of interest, it has been reported that depletion of Numb through a conditional knockout increased mitochondrial fragmentation in the kidney through Drp1 [15], a critical protein in mitochondrial fission. In contrast to results reported herein, in other studies of kidney mitochondria from cells lacking Numb, mitochondrial membrane potential and ATP generating capabilities were not altered [15]. Further studies are required to understand both the mechanisms and functional implications of the altered mitochondrial properties.

Surprisingly, knockouts of Numb/NumbL in muscle fibers resulted in only 6 shared gene expression changes. How these gene alterations contribute to the muscle weakness observed in Numb/NumbL cKO mice is unclear. It remains possible that some of these DEGs reflect a compensatory response of skeletal muscle to the consequences of Numb/NumbL cKO. Downregulated genes were *Sema7a* and *Tmem132a*, neither of which has been previously implicated in aging or muscle weakness. Sema7a is involved in immunity, axon guidance and integrin signaling [33] and may be linked to osteoporosis [34]. Tmem132a is one of five variants of the transmembrane 132 family of proteins. Little is known regarding its biology; of interest, it was recently shown to bind WLS, a Wnt ligand stabilizing protein. Tmem132a augments WLS-Wnt ligand interactions thereby increasing Wnt signaling [35]. Upregulated genes were *Itga4, Mib1* and *VezF1*. Itga4 (integrin A4 subunit) is a cell surface adhesion molecule; it is possible that upregulation of Itga4 may be a compensatory mechanism to stabilize structural integrity of skeletal muscle. VezF1 is a transcriptional regulator implicated in regulating angiogenesis; VezF1 knockdown has been linked to impaired cardiac muscle contractility without alteration of Ca^2+^ release kinetics [36]. We are not aware of prior studies of the function of VezF1 in skeletal muscle; elevation of VezF1 in Numb/NumbL cKO mice may be an adaptive response to overcome the deleterious effects on homeostasis or contractility of the knockouts. Mib1 (Mindbomb, E3 ubiquitin protein ligase) is known to positively regulate Notch signaling by ubiquitinating Notch receptors leading to their endocytosis. Recent data have shown that Mib1 is upregulated in skeletal muscle satellite cells by both androgen and estrogen dependent signaling and that absence of Mib1 in myofibers leads to depletion of satellite cells due to impaired cell cycle exit of proliferating satellite cells [37]. Mib1 has been suggested to ubiquitinate and target for proteasomal degradation other proteins such as the anti-apoptotic protein DAPK1, although implications for muscle fiber biology remain unclear.

An important control used in analysis of DEG induced by the Numb/NumbL cKO was analysis of effects of tamoxifen on gene expression in Numb^(f/f)^/NumbL^(f/f)^ mice which lack Cre. A noteworthy finding was that the number and list DEGs was quite different between male and female genotype controls. One possible explanation is the much higher levels of estrogens in female mice such that addition of a tamoxifen elicited fairly minimal effects on this already activated program of estrogen-regulated genes within skeletal muscle. Regardless, the findings argue for inclusion of genotype controls treated with tamoxifen or vehicles in RNA-sequencing studies of the effects of inducible knockouts on skeletal muscle gene expression patterns.

An unexpected finding was the reduced fusion index observed in myotubes formed from Numb depleted primary mouse myoblasts. The mechanisms underlying this observation are unclear. Some understanding of the roles of Numb in progenitor cells of the myogenic lineage have been gleaned using a conditional knockout under the Pax7 promoter [16]. These studies showed reduced ability of skeletal muscle to undergo repair after cardiotoxin injury associated with reduce proliferation of skeletal muscle satellite cells associated with upregulation of myostatin.

In summary, Numb protein levels decrease in skeletal muscle with age. Loss of Numb in skeletal muscle myofibers reduces contractile function, sarcomere length and mitochondrial size. Impaired release of calcium from intracellular stores may contribute to poor contractile function in the absence of Numb. Altered levels of LMs result from loss of Numb and may be key drivers of some of the physiological characteristics of Numb deficient muscle. Changes in levels of genes encoding an integrin (Itga4), a molecule involved in integrin signaling (Sema7a) and architectural changes in Z-lines suggest one common and potentially related structural theme in Numb deficient skeletal muscle fibers. Accumulation of residual bodies, small mitochondria, and prior evidence linking Numb to mitochondrial fission [15] further implicate Numb as having central roles in homeostasis of mitochondria in skeletal muscle and other tissues. A role for such perturbations in the well-described age-related decline in mitochondrial function is an attractive direction for future investigations.

## Supporting information

Supplementary Movie 1 (Numb cKO)

Supplementary Movie 2 (Control)

## Acknowledgements

We are grateful to Tom Rando and Rob Krauss for helpful discussions. The authors of this manuscript certify that they comply with the ethical guidelines for authorship and publishing in the Journal of Cachexia, Sarcopenia and Muscle [39].

## Data availability Statement

The authors declare that the data supporting the findings in this study are included in the paper and the Supplementary information or available upon request from the corresponding authors.

