## Supplementary Figures for "Numb is required for optimal contraction of skeletal muscle"

Address Correspondence to:

Christopher Cardozo, MD

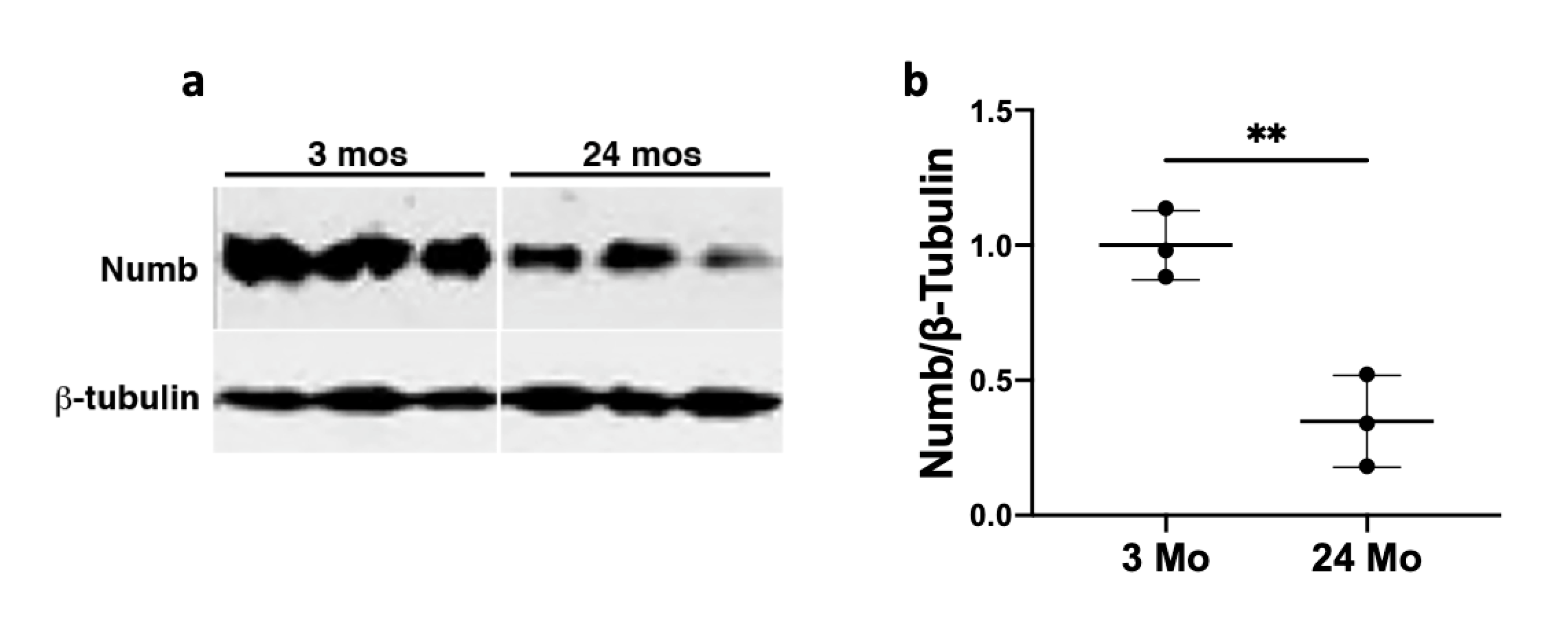

**Supplementary Fig. S1**. Expression of Numb protein in mouse muscle. Numb protein levels present in gastrocnemius muscles from 3 and 24-month-old C57BL/6N mice from the NIA Aged Rodent Colonies was detected by Western blotting and quantified using ß-tubulin as a loading control. (a) Non-contiguous lanes from a single representative western blot; (b) quantification of band intensities shown in a. where Numb protein levels were normalized using ß-tubulin. Data are presented as mean values ± STD. **, p < 0.01, t-test.

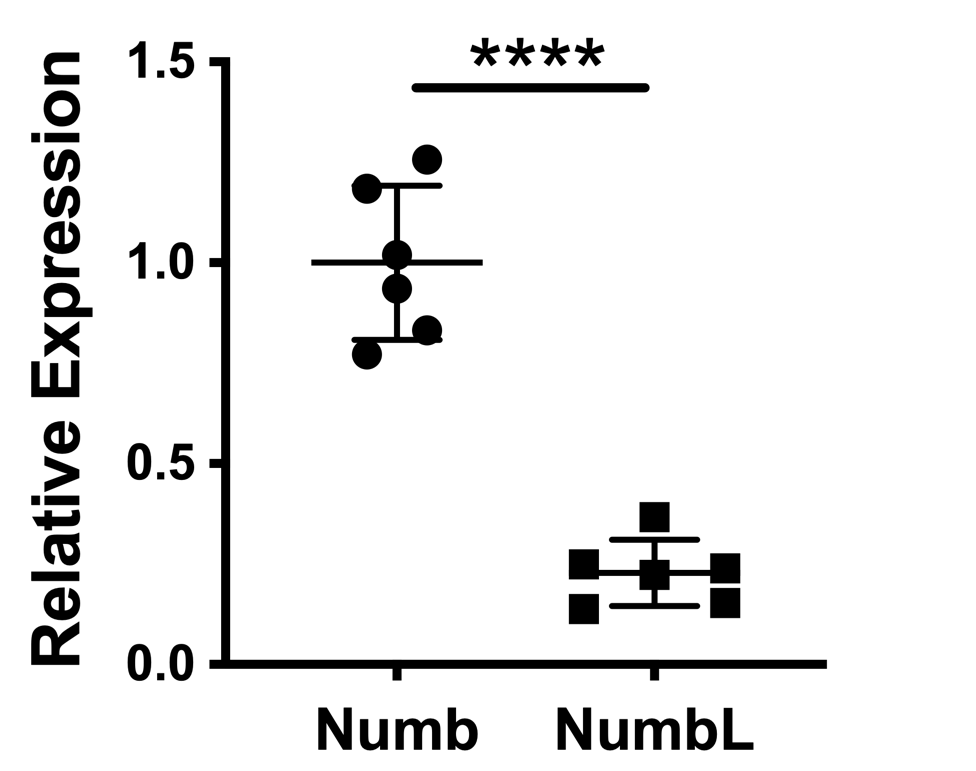

**Supplementary Fig. S2.** Expression of Numb and NumbL genes in mouse muscle. Total RNA was isolated from gastrocnemius muscle using RNeasy kits (Qiagen) and reverse transcribed using the High-Capacity cDNA reverse transcription kit. Numb and NumbL expression were determined by qPCR using specific TaqMan primers and probes and TaqMan universal Master mix (N=6). mRNA expression was calculated with the -2DDCt method using GAPDH as a housekeeping gene. NumbL expression was normalized relative to Numb. Statistical differences were analyzed by a two-tailed t-test; ****, p<0.0001. Data are expressed as mean values ± STD.

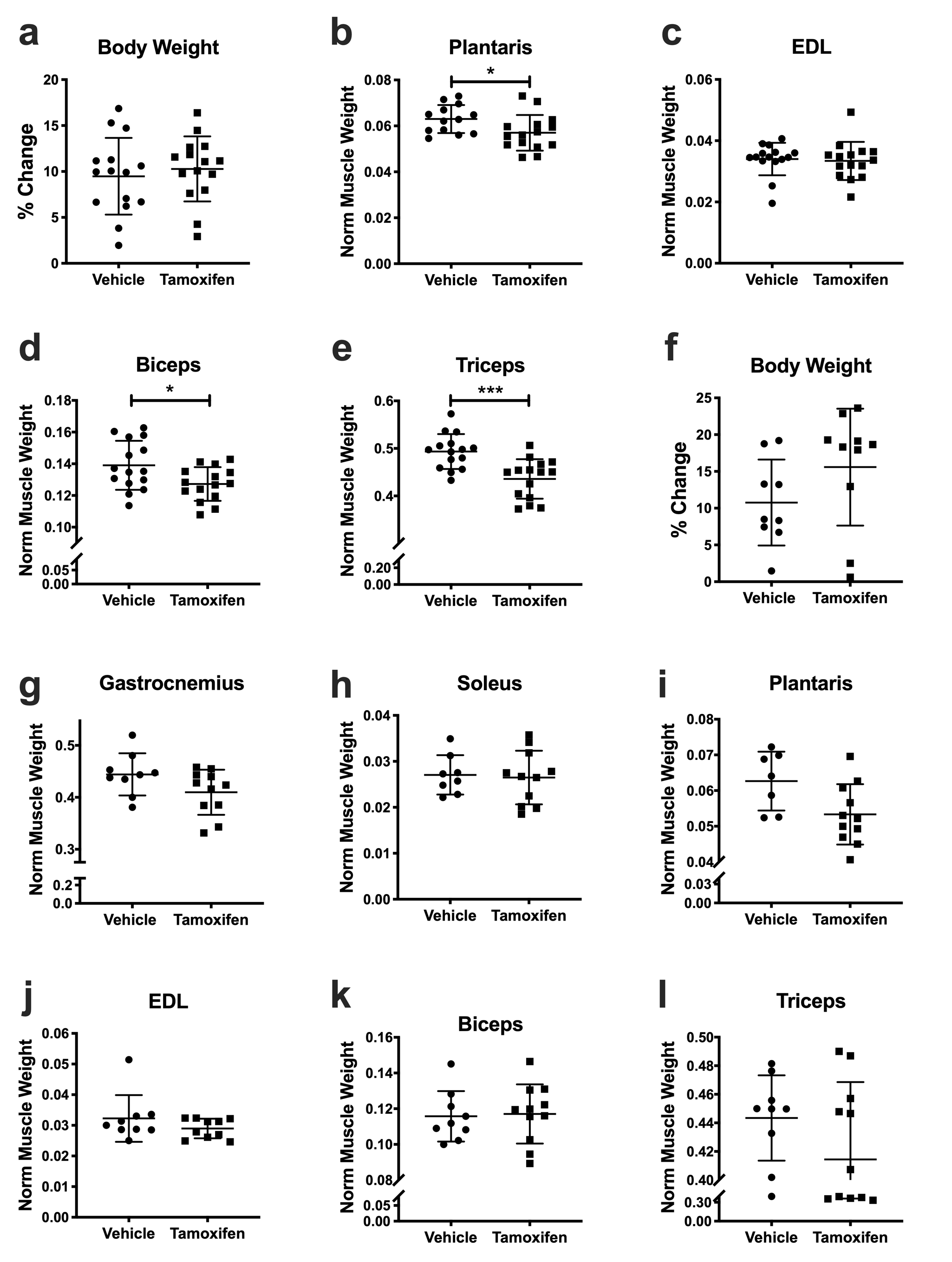

**Supplementary Fig. S3. Body and tissue weights.** **a-e:** data show body or muscle weights for HSA-MCM Numb^f/f^/NumbL^f/f^ mice at 56 days after the beginning of treatment with either tamoxifen or vehicle. **a,** Change in body weight (%) of vehicle or tamoxifen treated **b-e,** Normalized muscle weight of Plantaris (**b**), EDL (**c**), Biceps (**d**) and Triceps **(e)** (N=15/group).

**f-l:** data show body or muscle weights for Numb^f/f^/NumbL^f/f^ mice (genotype controls) at 56 days after the beginning of treatment with either tamoxifen or vehicle. **f,** Percent body weight change of vehicle or tamoxifen treated Numb^f/f^/NumbL^f/f^ mice 56 days after the beginning of the treatment. **g-j,** Normalized muscle weight of Gastrocnemius **(g),** Soleus (**h**), Plantaris (**i**), EDL **(l),** Biceps **(k)** and Triceps (**j**). (N=9 for vehicle-treated and N=11 for tamoxifen-treated animals). All muscle weights were normalized to pre-induction body weight. Statistical differences were analyzed by two-tailed t-tests; ***, p<0.0005, * p< 0.05.

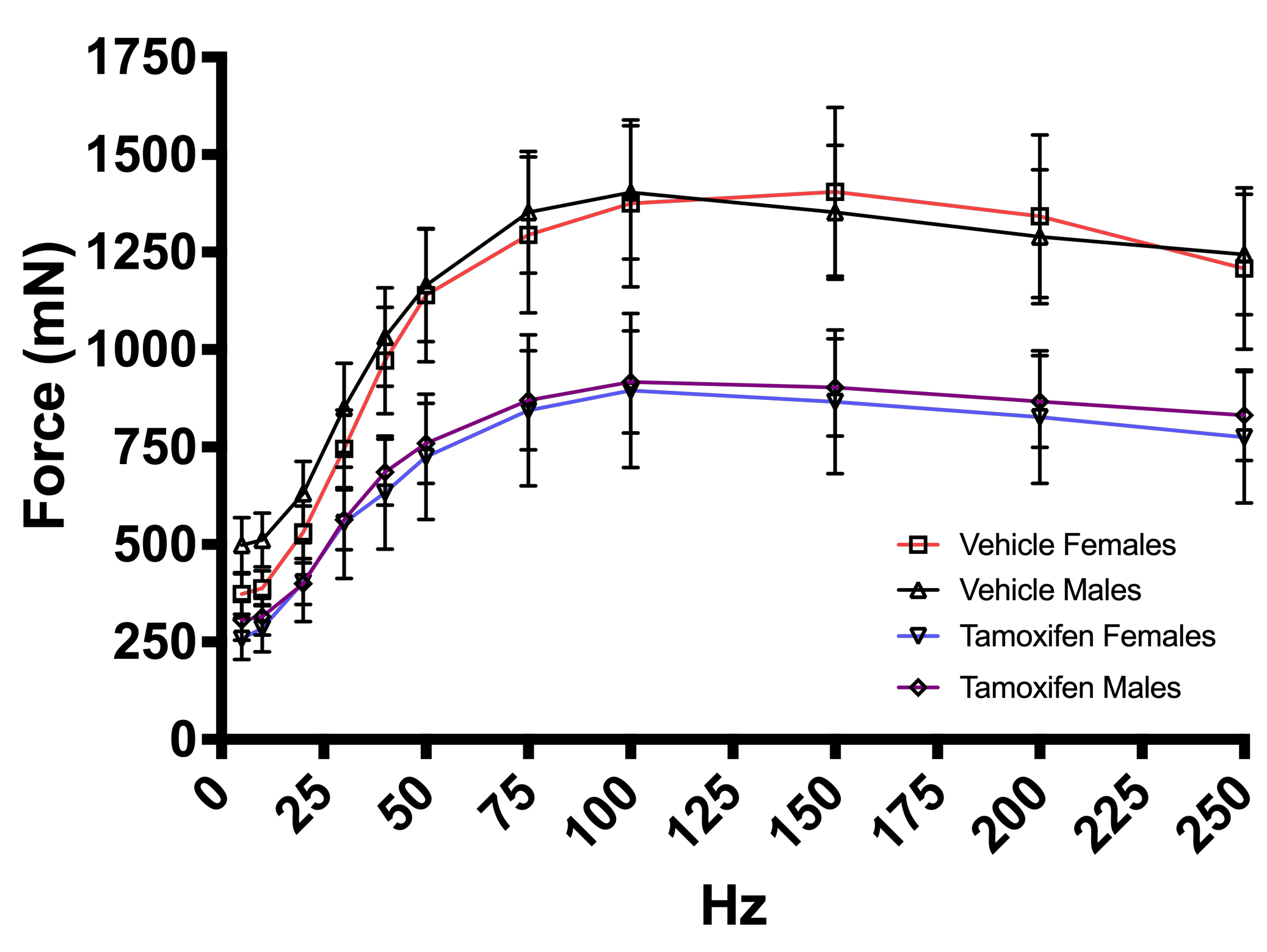

**Supplementary Fig. S4.** Force-frequency plots generated by *in-situ* physiological testing of gastrocnemius muscle from male or female mice at 56 days after beginning treatment with tamoxifen or vehicle. No difference was observed in force-frequency relationships between genders. Group sizes were: N = 8 for vehicle and tamoxifen treated females, N = 7 for vehicle treated males, N = 5 for tamoxifen treated males.

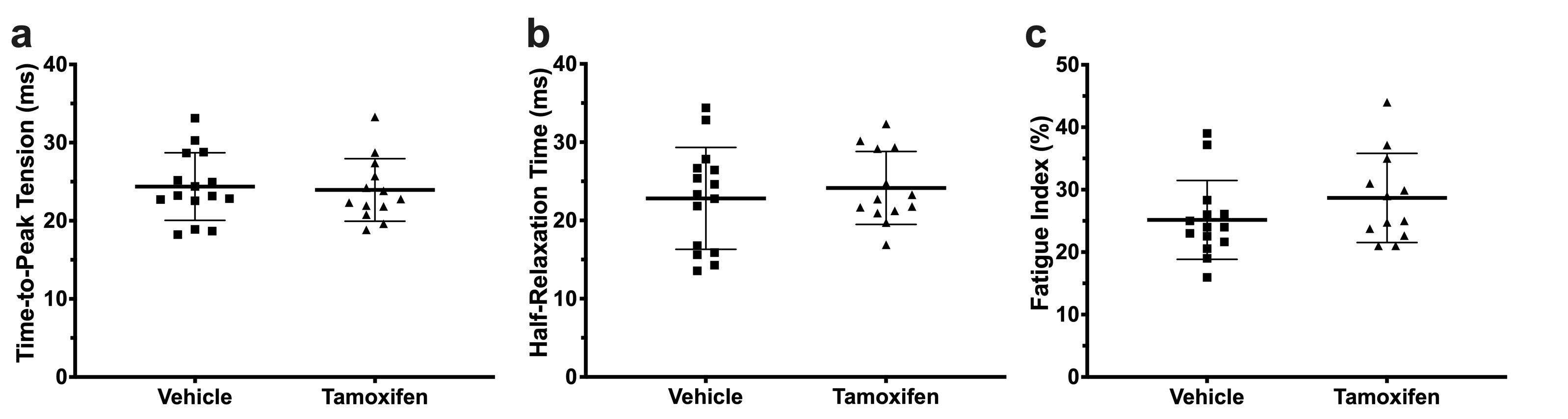

**Supplementary Figure S5**. *In-situ* physiology analysis of gastrocnemius muscle isolated from vehicle and tamoxifen treated HSA-MCM Numb^f/f^/NumbL^f/f^ mice 56 days after the beginning of the treatment. N=15 for the vehicle group and 13 for the tamoxifen group. **a,** Time to peak tension. **b,** Half relaxation time. **c,** Fatigue index. No statistically significant differences were found for these parameters (two tailed t-test p>0.05).

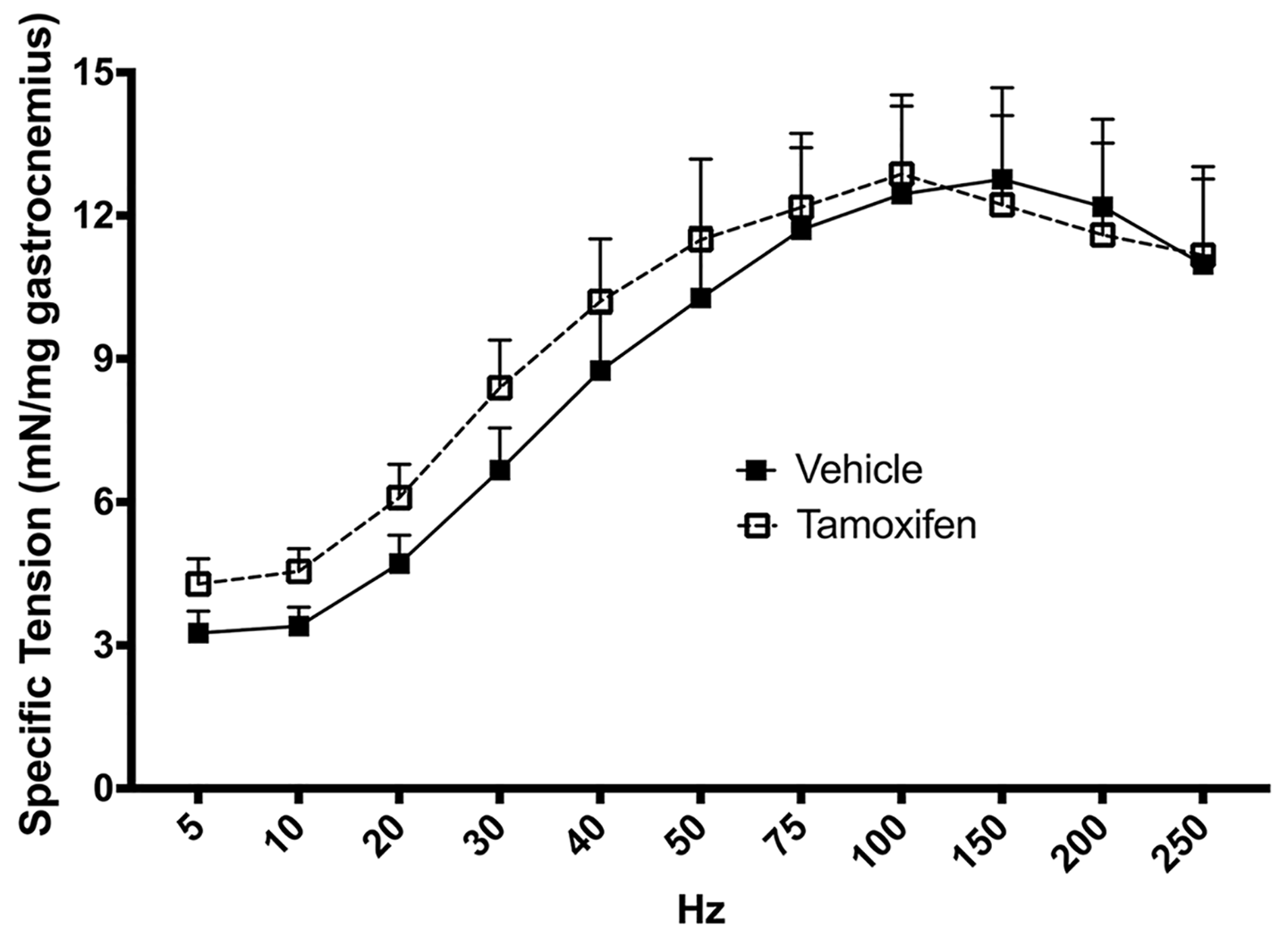

**Supplementary Fig. S6**. Specific tension-frequency curves determined by *in-*situ testing of gastrocnemius muscle in Numb^f/f^/NumbL^f/f^ mice (genotype controls) treated with tamoxifen or vehicle for 56 days after the beginning of the treatment. (N=8 animals for the vehicle group and N=5 for the tamoxifen group). Statistical analysis was performed with a repeated measure two way ANOVA . Standard error bars are displayed in the upward direction only to improve the ease with which data points can be seen in the figure. There was no significant interaction effect (F value = 0.5832, DFn 10, DFd 110, p = 0.8247 or main effect of tamoxifen (F value = 0.1479 with DFn 11, DFd 110, p=0.7079).

**Supplementary Fig. S7.** **Effects of Numb/NumbL cKO on fiber cross sectional area (CSA) and fiber type distribution.** Tibialis anterior (TA) muscle was isolated from HSA-MCM Numb^fl/fl^/NumbL^fl/fl^ mice treated with vehicle or tamoxifen. **a,** Representative hematoxylin-eosin stained cryosections. **b,** Mean fiber cross sectional area. **c,** Distribution of fiber cross sectional area in vehicle and tamoxifen treated TA. CSA was determined for 150 fibers on one image per animal for 3 animals per group. **d-e,** Representative immunostaining of muscle fiber types in TA from vehicle (**d**) or tamoxifen treated animals (**e**) **f-g,** Percent muscle fiber types in vehicle and tamoxifen treated animals. All fibers on a cross-section were scored; one section was analyzed for each of 3 animals per group. Scale bar: **a,** 400 μm; **d-e,** 25 μm. No significant differences were observed between vehicle and tamoxifen groups (two-tailed unpaired t-test).

**Supplementary Fig. S8.** Tibialis anterior (TA) muscle was isolated from Numb^fl/fl^/NumbL^fl/fl^ mice (genotype controls) treated with vehicle or tamoxifen for 56 days. **a,** Hematoxylin-eosin stained TA muscle from vehicle (left panel) and tamoxifen treated (right panel) animal. **b,** Mean fiber cross sectional area. **c,** Distribution of fiber cross sectional area (CSA). 150 fibers were scored/animal. Data are mean values ± SEM for 3 animals per group. **d**, Representative images of cryosections immunostained for detection of muscle fiber types in TA from vehicle (left panel) and tamoxifen treated animals (right panel) **e**, Percent muscle fiber types in vehicle and tamoxifen treated animals. A total of 2400 fibers were scored /animals (N=3/group). *<0.05, two-tailed unpaired t-test. Scale bar: **a**: 400 μm, **d**: 25 μm.

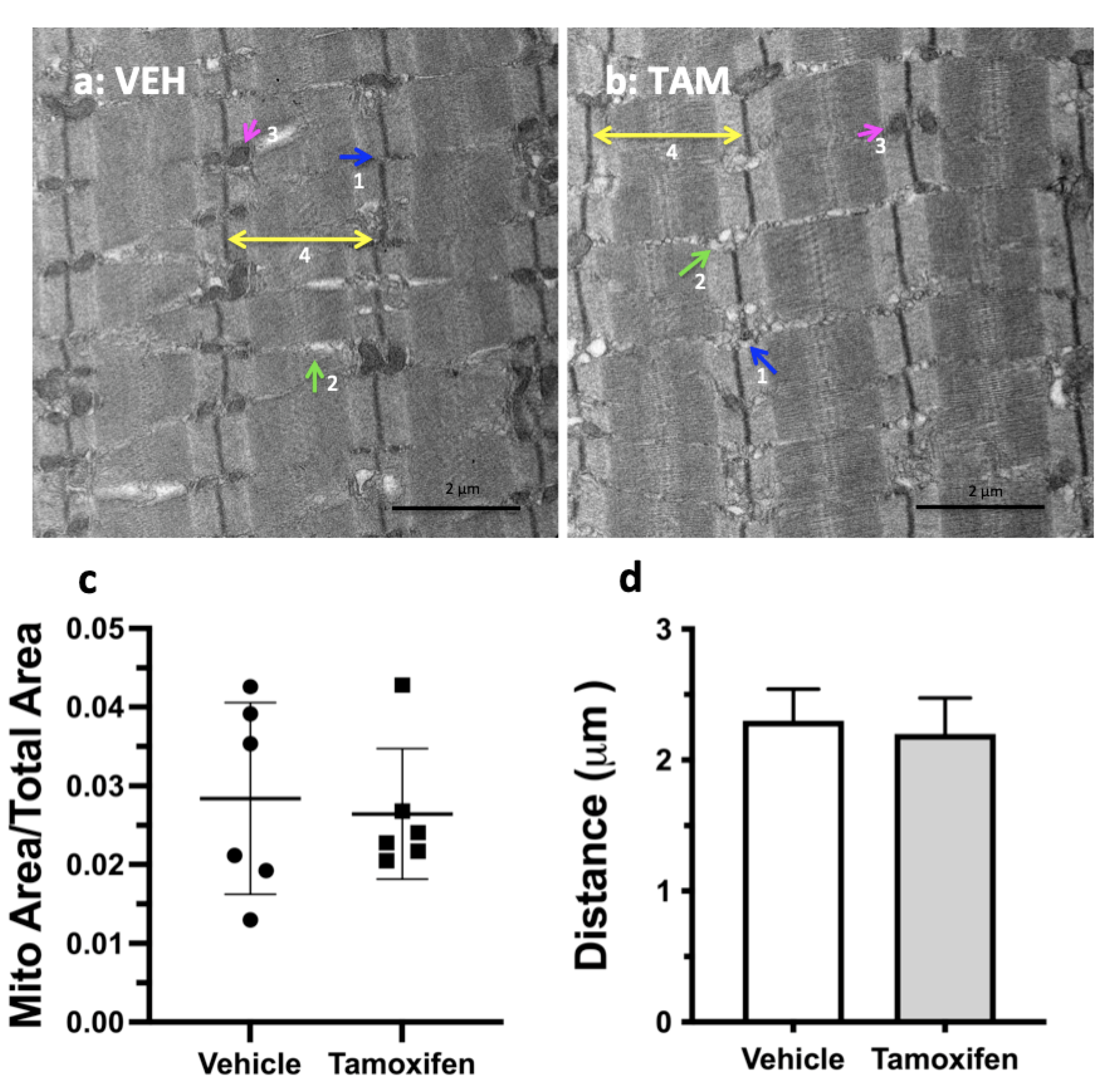

**Supplementary Fig. S9.** Representative transmission Electron Microscopy images acquired at 5800x of tibialis anterior muscle from genotype control (Numb(f/f)/NumbL(f/f)) mice are shown for Vehicle treated (**a**) and tamoxifen-treated (**b**) mice. Mitochondrial area as a fraction of total area was determined for 3 sections for each of 3 different mice for each gender (N=6 vehicle and 6 tamoxifen-treated mice) (**c)**. Mitochondrial and total area was determined using the Measure tool of NIH ImageJ. (**d**) Distance between Z-discs (μm) (n=3/per group). Tamoxifen treatment, tissue harvest and TEM were performed as described in Materials and Methods.

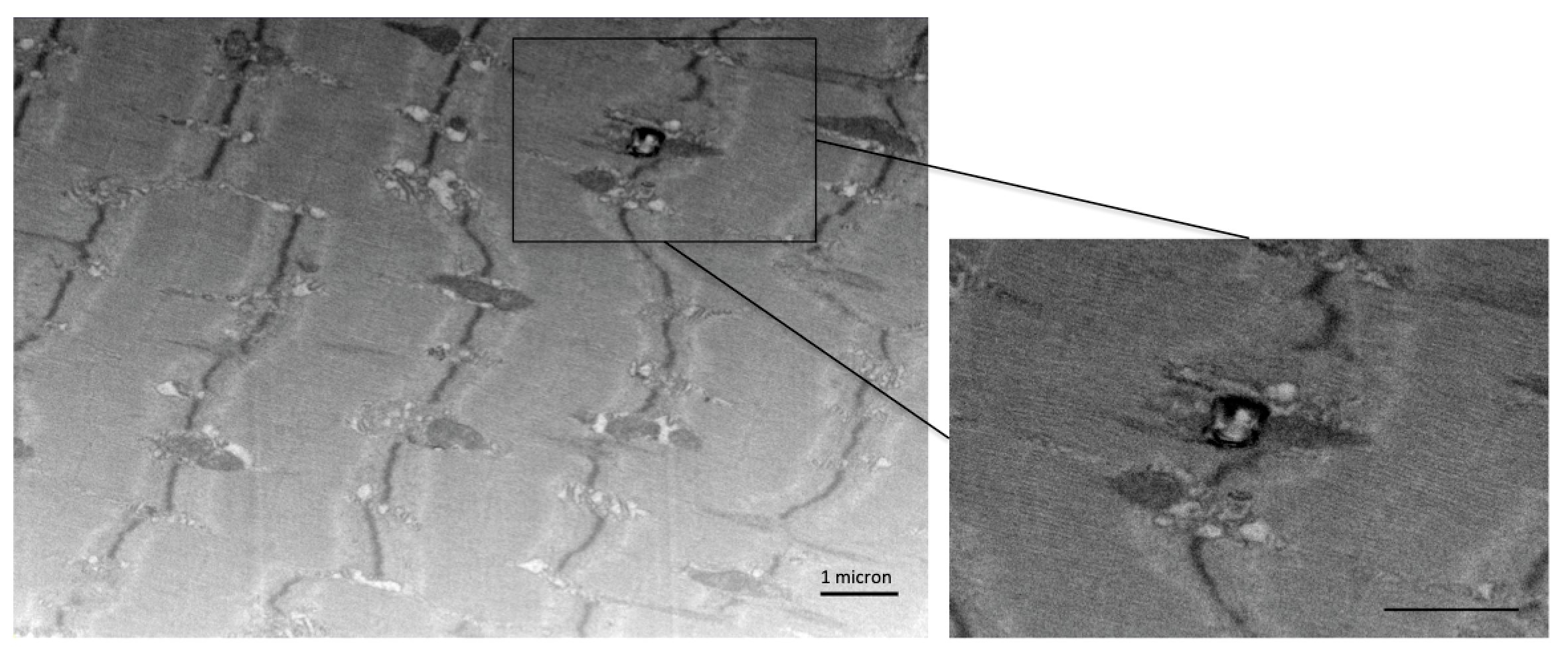

**Supplementary Fig. S10.** Representative FIB-SEM Image of tibialis anterior muscle from HSA-MCM/(Numb(f/f)/NumbL(f/f) mice treated with tamoxifen showing an example of residual bodies noted in the female Numb/NumbL cKO animals. The pop-out image shows an enlarged image of the area of interest delineated by a box in the original image. Tamoxifen treatment, tissue harvest and FIB-SEM were performed as described in Materials and Methods. The scale bar represents 1 micron.

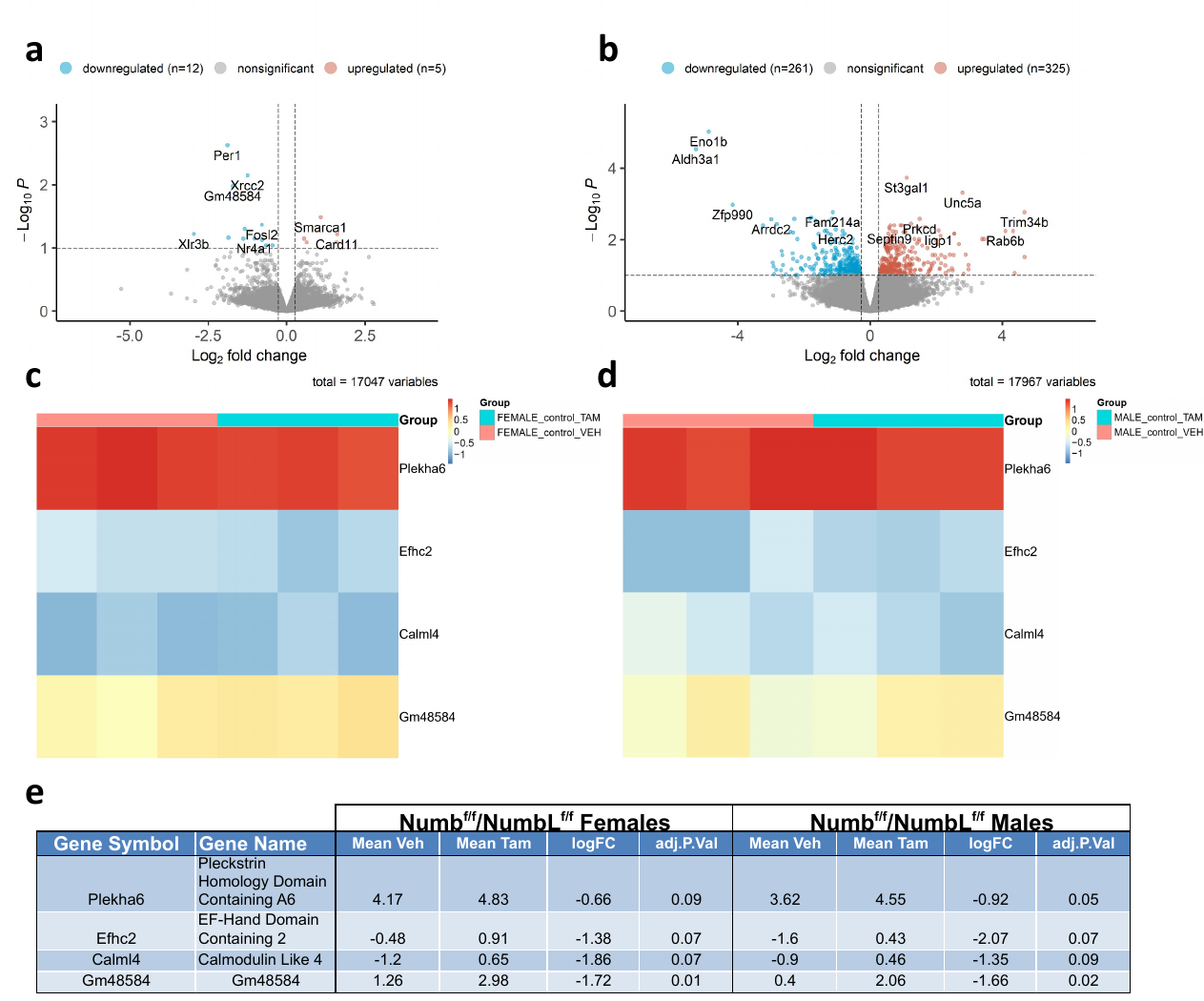

**Supplementary Fig. S11.** **RNA sequencing genes that are differentially expressed genes after tamoxifen treatment in genotype controls (Numb^(f/f)^/NumbL^(f/f)^ mice).** (**a**, **b**). Volcano plots showing mRNA expression levels for genes that had fold-changes > 1.2 with p values adjusted for FDR < 0. Light brown spots indicate upregulated genes, light blue spots downregulated genes. All other genes detected are shown as grey dots. Values for numbers of up and downregulated genes are show at the top right and top left of each panel, respectively. The total number of transcripts analyzed is shown under the X-Axis as Total Variables Heatmaps showing relative expression of genes altered in tamoxifen-treated females (**c**) or males (**d**) Numb(f/f)/NumbL(f/f) mice. (**e**) Table shows relative expression, log_2_ fold-change in expression and adjusted p value for the indicated genes for female (**c**) and male (**d**) mice.

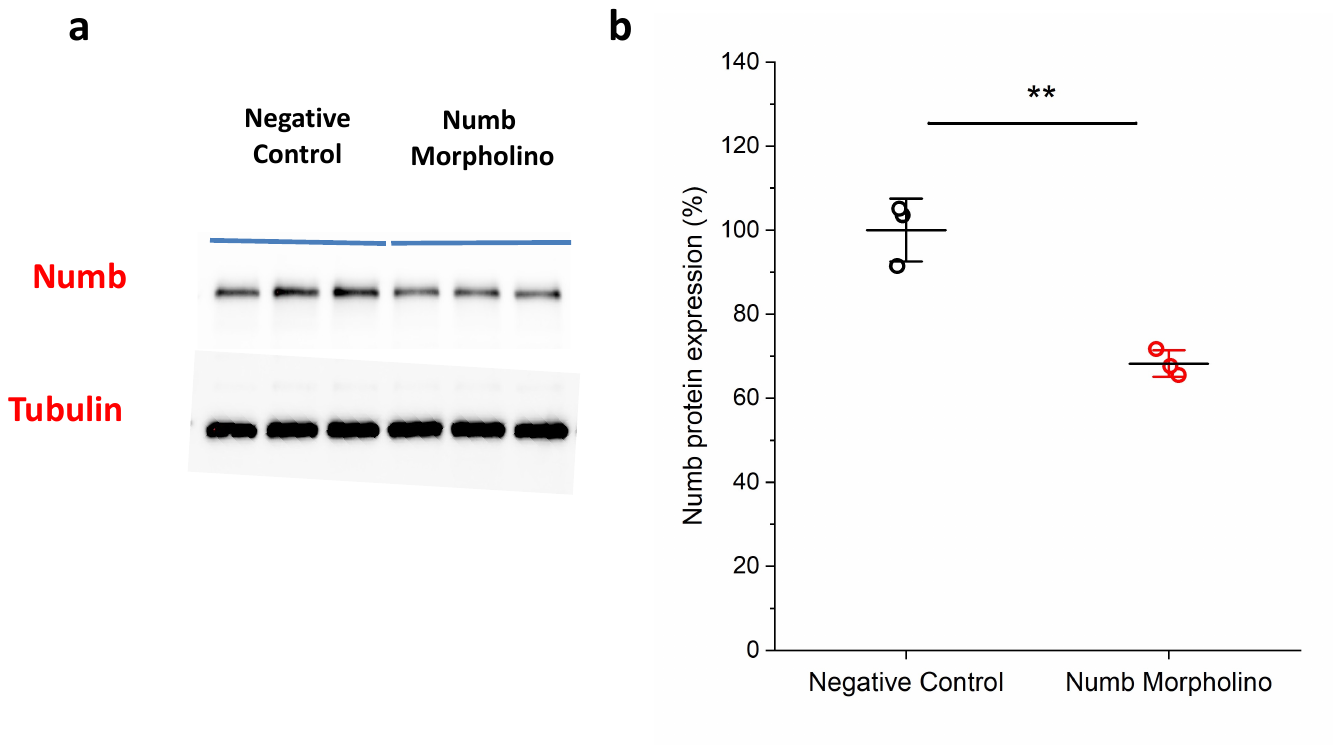

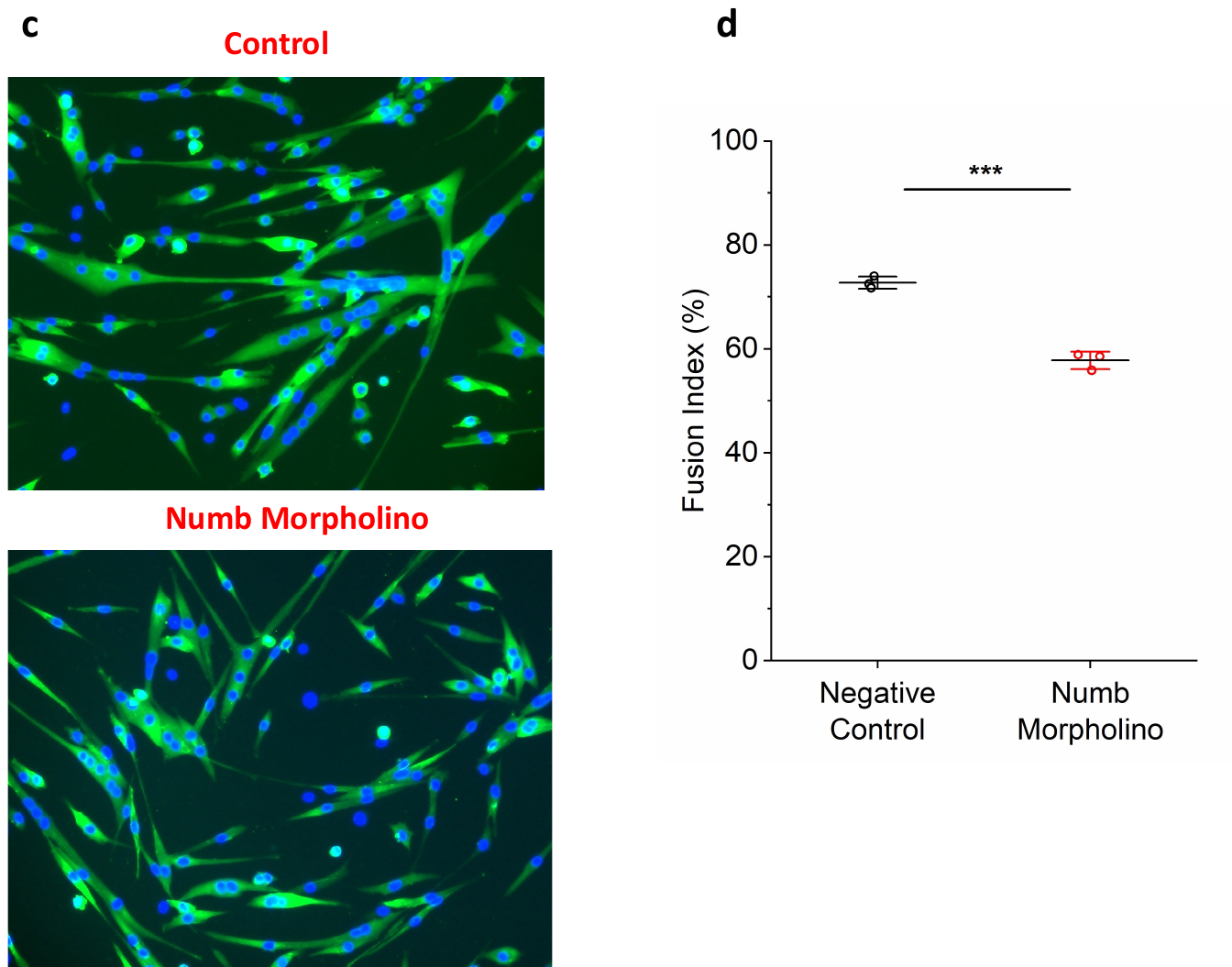

**Supplementary Fig. S12**. (a) Numb Western blot results after morpholino treatment for 48 h; (b) quantification of Numb Western blot results using ImageJ; (c**)** Downregulation of Numb significantly inhibit primary myoblast myogenic differentiation. (d) Treatment with a Numb morpholino significantly reduces fusion index. n = 3–4, **p < 0.01 and ***p < 0.001 compared with control, unpaired t-test.
