## Supplementary Methods for "Numb is required for optimal contraction of skeletal muscle"

Address Correspondence to:

Christopher Cardozo, MD

**Tissue harvest.** To harvest tissues, animals were anesthetized using inhaled isoflurane and muscle tissues were excised after careful dissection, weighed and immediately frozen in isopentane cooled in liquid nitrogen. Animals were euthanized by exsanguination under isofluorane anesthesia after completing sample collection.

**Immunofluorescence staining of tissue sections.** Frozen tissue was cut into 8 µm sections and mounted on glass slides (Fisher*brand* 12-550-15). The slides were dried for 30 min under a gentile stream of air and then fixed with freshly made, ice cold 4% paraformaldehyde (PFA) in PBS for 8 min with gentle shaking. Slides were than rinsed three times in PBS. The sections were permeabilized and blocked in PBS, 15% normal goat serum, 0.3% Triton X-100 (blocking solution), for 60-120 minutes at room temperature then incubated for 16-48 h at 4°C with anti-Numb (#2756, 1:50, Cell Signaling Technology,) and anti-dystrophin (ab3149, 1:100, Abcam) antibodies diluted in the above blocking solution. Slides were rinsed in PBS 3 times for 2 min each and the appropriate secondary antibodies (Alexa Fluor 647 conjugated anti-rabbit IgG F(ab’)_2_ #4414 and Alexa Fluor 488 conjugated anti-mouse IgG F(ab’)_2_ #4408, both from Cell Signaling) were applied at 1:200 in blocking solution for 60 min at room temperature or overnight at 4°C. Slides were rinsed in PBS, 0.1% Triton X-100, 2 times for 10 min each and again in PBS for 5 min. The sections were mounted using Vectashield mounting media with 4’,6 diamidino-2-phenylindole (DAPI) (Vector Laboratories) and imaged with a Zeiss LSM700 confocal microscope. Images were acquired with the Zeiss Zen software and processed with Adobe Photoshop CS.

**Fiber type composition and cross-sectional area measurement.** Determinations of fiber type distribution and cross-sectional area were performed on sections of TA muscle isolated from HSA-MCM Numb^f/f^/Numb^f/f^ mice treated with vehicle or tamoxifen and harvested at 56 days post-induction (n=3/group). For fiber typing, frozen TA muscle was cut in 10 μm sections with a cryostat, fixed in ice-cold 4% PFA in PBS for 10 min and permeabilized and blocked as above for 2 h. The sections were first incubated with anti-myosin heavy chain IIA (MHC IIA) antibody A4.74-c (1:50, Developmental Hybridoma Bank) overnight at 4^o^C and the signal revealed using the AP-conjugate and reagents in the Expose Mouse and Rabbit specific AP detection IHC kit (Abcam). The tissue was then rinsed with PBS and incubated with anti-Myosin heavy chain IIB antibody 10F5 (1:5, Developmental Hybridoma Bank) overnight at 4^o^C and the signal revealed with Expose Mouse and Rabbit Specific HRP/DAB Detection IHC Kit (Abcam). The slides were mounted using Cytoseal XYL, cover slipped, and imaged using a Nikon Eclipse E-600. The images were processed with Abobe Photoshop CS. One section was scored per muscle for each of 3 animals per group. On average 2400 fibers were scored/animal.

For determination of cross sectional area, frozen TA was cut in 12 µm sections using a cryostat and stained with the Rapid-Chrome Hematoxylin Eosin (H&E) Frozen Section Staining Kit (ThermoFisher). Tiled images were taken at 20x magnification with an EVOS FL Auto Cell Imaging System (Life Technology) and similar areas were identified for fiber analysis. The cross-sectional area of 150 fibers was measured using ImageJ software.

### Isolation, Culture and Immunofluorescence staining of muscle fibers. Muscle fibers were isolated from mixed hindlimb muscle essentially as described [1]. The explanted muscles were digested with 400 U /ml of collagenase type I (Worthington) in DMEM-F12 medium at 37 ^0^C for 55 min. Fibers were collected with a fire-polished pipette in DMEM-F12 supplemented with 15% horse serum and antibiotic-antimycotic solution (Life Technology). To remove debris, the fibers were re-picked 2-3 times in fresh medium. The fibers were either fixed immediately with 4% paraformaldehyde in PBS for 10 minutes or cultured in the above medium for 48-72 h and then fixed as above.

Free floating fibers were blocked for 1 h in Tris buffer saline (TBS) containing 10% normal goat serum and 0.3% Triton X-100 followed by overnight incubation at room temperature with primary antibodies diluted in the above blocking solution. After washing with PBS, the fibers were incubated with Alexa Fluor^®^-488 conjugated anti-rabbit IgG and Alexa Fluor^®^ 568 conjugated anti-mouse IgG antibodies (1:400 dilution in blocking solution, ThermoFisher) for 2 h and washed with PBS. Nuclei were stained with DAPI (1 μg/μl in PBS) and the fibers mounted on slides with Fluorogel mounting medium (Electron Microscopy Sciences). The following primary antibodies were used for staining: anti-Numb (1:300, Cell Signaling Technology), anti-dystrophin (1:100, ab3149, Abcam)**,** anti-actinin (1:300, ab9465, Abcam) and anti-dihydropyridine receptor (DHPR) (1:100, mAb 23-920, ThermoFisher). The stained fibers were imaged with a Zeiss LSM700 confocal microscope. Images were acquired with the Zeiss Zen software and processed with Adobe Photoshop CS.

**Western blotting**.

For Western blotting of tissues, tissue was homogenized in 10 mM Na phosphate pH 7.4, 150 mM NaCl, 2 mM EDTA, 1% Triton X-100, 0.5% Na^+^ deoxycholate, 1% SDS supplemented with Halt protease and phosphatase inhibitor cocktail (ThermoFisher). The lysates were centrifuged at 14,000 rpm for 15 min and the supernatant saved. Protein concentration was determined using the BCA reagent (Pierce). Proteins (50 μg) were separated by 4-15 % SDS- PAGE gels and transferred onto Immobilon P (EMD Millipore) membranes. The membranes were blocked in Tris buffer saline, 0.5% Tween-20 (TBST), 5% non-fat dry milk and incubated overnight at 4°C with rabbit monoclonal anti-Numb ((#2756, Cell Signaling Technology) 1:1000 in blocking solution). The membrane was incubated with peroxidase conjugated anti-rabbit IgG (GE Healthcare) (1:7500 in blocking solution) and the bands revealed by incubation with ECL Prime reagent (GE Healthcare) and captured using to film or an Imager 600 gel scanner (GE Healthcare). The blots were stripped and probed with anti-GAPDH (sc-365062, 1:2,000, Santa Cruz) as loading control. Bands were quantitated with ImageQuant (GE Healthcare). Numb levels were normalized to GAPDH and expressed relative to the average of sham levels.

For Western blotting of mouse primary myoblast samples, ~30 µg of total proteins were fractionated by 4–15% Mini Protean TGX gels (Bio-Rad) and transferred to polyvinylidene difluoride (PVDF) membranes (Bio-Rad). Membranes were blocked in 5% non-fat dry milk in TBST for 1 h at room temperature (RT), followed by incubation with rabbit monoclonal anti-Numb (Cell Signaling Technology) or β-tubulin (1:1000, Cell Signaling Technology) antibody in 5% bovine serum albumin (Millipore Sigma) in TBST at 4 °C overnight. HRP-conjugated goat anti-rabbit (1:10,000, Jackson ImmunoResearch) secondary antibody was then applied to membranes for 1 h at RT. After five 5-min washes in TBST, Clarity Max ECL Western blotting substrates (Bio-Rad) were used to detect the signal using ChemiDoc MP imaging system (Bio-Rad). Images were quantified with ImageJ.

***In-situ* physiology**

*In* *vivo* stimulations, data collection and data analysis of gastrocnemius muscle were carried out using an Aurora Scientific 310C Complete In Vivo Physiology System (Aurora Scientific; Toronto, Canada). Force values were captured with the Aurora Scientific DMC acquisition software and analyzed with the Aurora Scientific DMA analysis software. Physiological testing was done as described [2] with minor modifications. All surgeries and testing occurred with the animals under anesthesia achieved by inhalation of 2-5% isoflurane and placed on a heating block. The hair was removed from the upper portion of the left hindlimb and the skin cleaned with 70% ethanol. The sciatic nerve was exposed using an incision and blunt dissection and the skin of the lower limb was removed to expose the gastrocnemius/plantaris/soleus complex. After removing the plantaris and soleus, the gastrocnemius was isolated by cutting the calcaneus to free a portion of the calcaneus with the Achilles tendon still attached; the distal end of the gastrocnemius was tied to a length-force transducer using a silk suture around the Achille’s tendon just proximal to the bone fragment. The knee was fixed into place with a pin vise and the sciatic nerve was connected to an external digital stimulator using a cuff electrode. The muscle and nerve were kept warm using a surgical lamp and kept moist by bathing them with warmed (37˚ C) lactated Ringer’s solution. Variables determined were peak isometric twitch (P_t_) and tetanic (P_o_) force as well as a force-frequency distribution, time-to-peak tension (TPT), half-relaxation time (HRT) and the fatigue index (FI). Optimal muscle length (L_o_) was determined by 20 ms, 1 V square-wave pulse twitch contractions and increasing the muscle length by 0.5 mm until force dropped or failed to increase. One min rest intervals took place after each twitch contraction; a piece of stiff suture was cut to the respective L_o_ and the muscle was measured against it after every contraction and returned to proper length when needed. Once L_o_ was determined there was a 2 min rest interval after which P_t_, TPT and HRT were determined using the average of 10 supramaximally-stimulated twitch contractions (200 ms, 4-5 V square-wave pulse). After the last twitch contraction, there was 2 min of rest and the tetanic contractile force was determined by a force-frequency relationship in which the muscle was stimulated with trains of electrical stimulation (4-5 volts, 1ms pulse duration and train duration of 300 ms) at 5, 10, 20, 30, 40, 50, 75, 100, 150, 200 and 250 Hz with 2 min of rest between each contraction between 5 and 50 Hz and 5 min of rest between 75 and 250 Hz. There was 5 min of rest after the last tetanic contraction and the FI was measured using 120 consecutive isometric contractions using supramaximal stimulations at 40 Hz for 300 ms followed by 1 s of rest. FI was calculated as the force of the final contraction divided by the contraction with the highest force.

**Transmission Electron Microscopy analysis.**

Mice were perfused with 2.5% glutaraldehyde, 2% paraformaldehyde in 0.1 M sodium cacodylate buffer, pH 7.4 (EMS). Muscle tissue was removed after careful dissection, cut in small pieces and post-fixed in the above solution for 24h at 4°C. The samples were treated with 1% osmium tetroxide in 100 mM cacodylate buffer pH 7.4 for 1 h, washed in distilled water four times (10 min/wash), and then treated with 2% aqueous uranyl acetate overnight at 4°C in the dark. Samples were washed and sequentially dehydrated with increasing concentrations of ethanol (20, 30, 50, 70, 90, and 100%) for 30 min each, followed by three additional treatments with 100% ethanol for 20 min each. Samples were then infiltrated with increasing concentrations of Spurr’s resin (25% for 1 h, 50% for 1 h, 75% for 1 h, 100% for 1 h, 100% overnight at room temperature), and incubated overnight at 70°C in a resin mold. Sections of 80 nm were cut on a Leica ultramicrotome with a diamond knife. Imaging then took place using a Talos L120C transmission electron (ThermoFisher) microscope operating at 120 kV. Images were acquired and analyzed with the TIA (TEM Imaging and Analysis) software.

Distance between Z lines was measured with Image J using 3-5 individual EM images per animal at a magnification of 5,300x for a total of 3 animals/group. 10-12 individual sarcomeres were measured per image. Data are presented as mean and standard deviation of Z-lines distances in each group. The arbitrary length units were converted to microns using the scale bar.

**Focused Ion Beam Scanning Electron Microscopy.**

The sample block was trimmed then mounted on a SEM sample holder using double-sided carbon tape (EMS). Colloidal silver paint (EMS) was used to electrically ground the exposed edges of the tissue block. The entire surface of the specimen was then sputter coated with a thin layer of gold/palladium. The tissue was imaged using back scattered electron (BSE) mode in a FEI Helios Nanolab 650. Images were recorded after each round of ion beam milling using the SEM beam at 2.0 keV and 200 pA with a working distance of 2.5 mm. Data acquisition occurred in an automated way using the Auto Slice and View G3 software. The raw images were 3,072 2,048 px2, with 20 nm slices viewed at a -38 degree cross-sectional angle. Each raw image had a horizontal field width (HFW) of 11.98 µm with an XY pixel size of 3-5 nm and 20 nm Z step size. These images were then aligned using the image processing programs in IMOD [3].

**Three-dimensional Reconstruction of Mitochondria and Quantitation.**

Serial sections of FIB-SEM images were reconstructed into three-dimensional representations of individual intermyofibrillar mitochondria using the computer visualization program *IMOD* *ver* 4.9 [3]. Intermyofibrillar mitochondria were chosen based on presence of clear borders. Mitochondria were manually traced every ~3 sections and then the boundaries of mitochondria were computationally estimated by interpolation and manually verified and adjusted as required. The total meshed volumes (nm^3^) and maximum lengths of each of the mitochondria (nm) were measured using the imodinfo and measure tools of IMOD, respectively. Movies showing locations in the image stack of the mitochondria that were reconstructed, and 3-dimentional models of those mitochondria as they are rotated in space, were generated in IMOD.

**RNA-Sequencing**.

Total RNA was isolated from gastrocnemius muscle using TRIzol followed by further purification using RNeasy Kits (Qiagen). RNA sequencing was performed at the NYU Genome Technology Center. Sequences were aligned to GRCm38 (mm10) mouse genome using Spliced Transcripts Alignment to a Reference (STAR) software and gene-level read counts were generated by FeatureCounts. Genes with expression of 0 in more than 30% of all samples were removed; data were normalized using limma/voom. We performed differential expression analysis using the c package v3.44.3 from Bioconductor [4]. Benjamini-Hochberg adjusted P values of less than or equal to 0.1 and fold change greater than or equal to 1.2 were used as the thresholds for detecting DEGs. Volcano plots were generated using the EnhancedVolcano package v1.6.0 from Bioconductor.

**Muscle Lipidomic Profiling**

All components of LC-MS/MS system are from Shimadzu Scientific Instruments, Inc. (Columbia, MD). LC system was equipped with four pumps (Pump A/B: LC-30AD, Pump C/D: LC-20AD XR), a SIL-30AC autosampler, and a CTO-30A column oven containing a 2-channel six-port switching valve. The LC separation was conducted on a C8 column (Ultra C8, 150 × 2.1 mm, 3 µm, RESTEK, Manchaca, TX). The MS/MS analysis was performed on Shimadzu LCMS-8050 triple quadrupole mass spectrometer. The instrument was operated and optimized under both positive and negative electrospray and multiple reaction monitoring modes (+/− ESI MRM). The settings of flow rate and gradient program for the LC system as well as MS/MS conditions are recommended by a software method package for 158 lipid mediators (Shimadzu Scientific Instruments, Inc., Columbia, MD) and further optimized following previously published quantification method [5]. The *m/z* transitions (precursor to product ions) and their tuning voltages were selected based on the best MRM responses from instrumental method optimization software. All analyses and data processing were completed on Shimadzu LabSolutions V5.91 software (Shimadzu Scientific Instruments, Inc., Columbia, MD).

**Tissue Preparation for Lipidomic Analyses**

Gastrocnemius muscles were isolated after mice sacrifice, then snap frozen in liquid nitrogen immediately and stored at -80°C. Before the experiment, the aliquoted frozen muscle tissue (50-100 mg) was defrosted on ice, weighed carefully, and minced into small pieces. The minced muscle was added into 1.0 mL ice-cold 80% (v/v) methanol in water, then was homogenized using a TissueLyser II homogenizer (Qiagen, Germantown, MD) at the frequency of 30 s^-1^, in 8×30-s bursts, waiting 20 s in between to avoid high temperature. The obtained homogenate was mixed with 5 µL of isotope-labelled LM internal standards (IS) mixture stock solution (5 µg/mL for AA-d_8_, 2 µg/mL for DHA-d_5_ and EPA-d_5_, and 0.5 µg/mL for the rest IS), and then agitated on ice and in the dark for 1-2 h, followed by centrifugation at 16,000 ×g at 4°C for 10 min to remove any tissue residue and precipitated proteins.

Samples were cleaned and concentrated by Solid Phase Extraction (SPE) before being injected into the LC-MS/MS. Ice-cold 0.1% (v/v) formic acid (4 mL) was added to the obtained supernatant to fully protonate the LM species before sample was loaded onto preconditioned SPE cartridges (Strata-X 33 µm polymeric reversed phase, Phenomenex, Torrance, CA). Cartridges were washed with 0.1% (v/v) formic acid in water followed by 15% (v/v) ethanol in water to remove excess salts. Then the LMs from the SPE sorbent bed were eluted by methanol. Solvents were removed using an Eppendorf^®^ 5301 concentrator centrifugal evaporator (Eppendorf, Hauppauge, NY). The dried extracts were stored at -80°C for future LC-MS/MS analysis.

**Isolation, culture and myogenic differentiation of mouse primary myoblasts**

Mouse primary myoblasts were prepared from hindlimb muscle by digestion with 0.1% pronase (Worthington) and were maintained/expanded in collagen I (R&D Systems) coated flasks in growth medium (GM) consisted of Ham F10, 20% feral bovine serum, 100 μg/ml streptomycin and 100 U/ml penicillin (ThermoFisher) supplemented with 5 ng/ml basic recombinant human fibroblast growth factor (Promega) as described [6]. For differentiation, culture medium was switched to differentiation medium (DM) containing high-glucose DMEM, 2.5% horse serum (Hyclone Laboratories), 100 U/mL penicillin, and 100 μg/mL streptomycin.

**Quantification of myogenic differentiation using fusion index**

Purified myoblasts were plated on E-C-L (MilliporeSigma)-coated 6-well plates at ~200,000 cells/well and cultured in DM for 48 h with 1 µM control or Numb morpholino (gene tools) treatment. After differentiation, cells were fixed in 10% neutral buffered formalin solution (NBF, MilliporeSigma) for 15 min. After removal of NBF, cells were washed 4 times with PBS, followed by permeabilization with 0.1% Triton X-100 in PBS for 15 min. Cells were then incubated with myosin heavy chain (MHC) fluorescein-conjugated antibody (1:100, FAB 6118A, R&D Systems) overnight at 4 °C. After 3 washes with PBS, DAPI (1:1000, MilliporeSigma) was added for 10 min incubation at room temperature. Images were taken with Olympus IX50 system using software cellSens Dimension 1.15 (Olympus Corp.). To quantify myogenic differentiation, fusion Index (FI) was calculated. FI was defined as: percentage of nuclei within myosin heavy chain‐expressing myotubes. Three independent experiments with two replicates in each experiment were performed.

**Mitochondrial function measurement**

Mitochondrial functions were determined using Seahorse XFe24 Analyzer (Agilent Technologies). Mouse primary myoblasts were cultured in collagen I (R&D Systems)-coated 6-well plates at ~150,000 cells/well in GM, followed by treatment with 1 µM control or Numb morpholino (gene tools) for 48 h. Cells were then collected and plated in Seahorse XFe24 cell culture microplates (Agilent Technologies) coated with E-C-L (MilliporeSigma) in DM for 48 h. Mitochondrial functions were evaluated using Seahorse cell energy phenotype test kit (Agilent Technologies) and Seahorse cell mito stress test kit (Agilent Technologies) according to instructions from the manufacturer. Four independent experiments with eight repeats in each experiment were performed.

**Intracellular calcium homeostasis**.

For measuring intracellular calcium homeostasis, purified mouse primary myoblasts were plated in E-C-L cell attachment matrix (Millipore Sigma) coated glass bottom dishes. After culture in differentiation medium (DMEM with 2.5% horse serum, 100 μg/ml streptomycin and 100 U/ml penicillin) for 12 h, myoblasts were treated with 5 nM vivo-morpholino oligo targeted to Numb (5’AGCTTTGCCGTAGTTTGTTCATGTT) or standard vivo-control oligo (both from Gene Tools). Differentiation was then continued for another 48 h for myotube development. The intracellular calcium transients produced by the stimulation of calcium release from SR with 20mM caffeine were measured using a Photon Technology International (PTI) imaging system. The differentiated myotubes were loaded with 2 µM Fura-2 AM (a radiometric calcium dye) and imaged in real time with the 14-BIT CoolSNAP CCD camera. All calcium imaging was analyzed with PTI EasyRatioPro fluorescence imaging software. Three to four independent experiments were conducted, resulting in 20-30 myotubes analyzed per group [7].
