## Supplementary figures and images for "Numb is required for optimal contraction of skeletal muscle"

### Supplementary Table 5

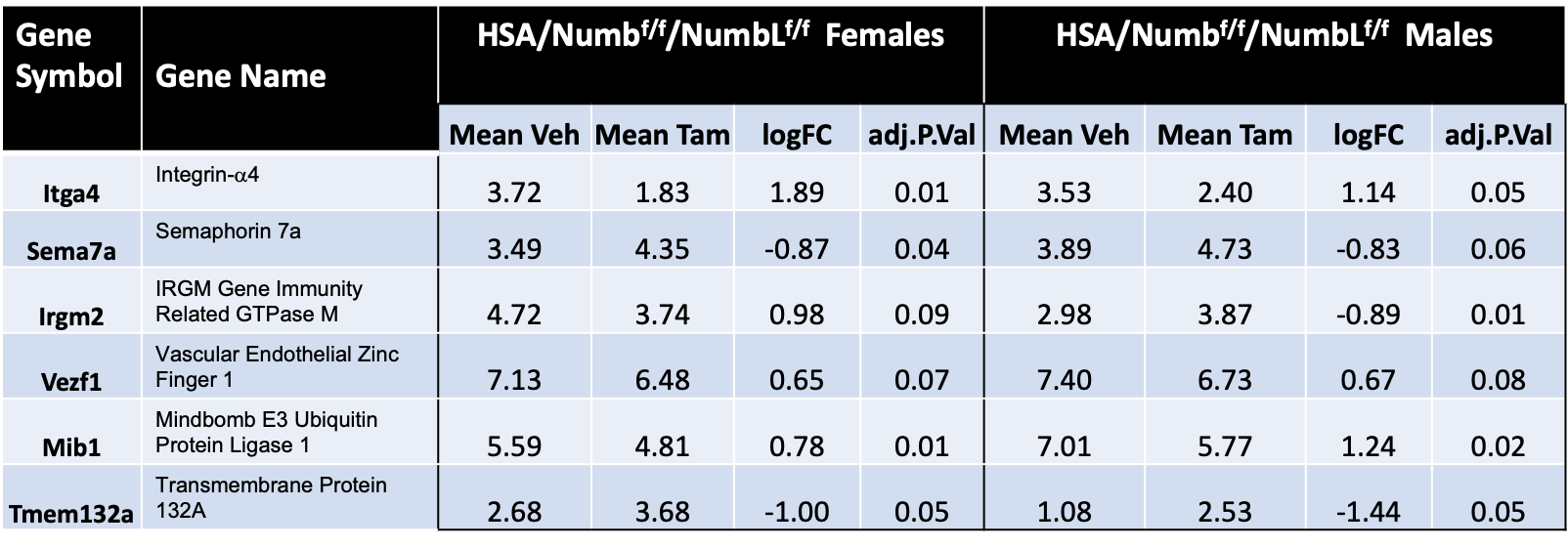
